# Eyes on the prize: a neural basis of perceptual choking under pressure

**DOI:** 10.64898/2026.09.30.755534

**Authors:** Megan E. McDonnell, Adithya Narayan Chandrasekaran, Chris Ki, Adam L. Smoulder, Hiroo Miyata, Byron M. Yu, Aaron P. Batista, Matthew A. Smith, Steven M. Chase

**Affiliations:** Neuroscience Institute, Carnegie Mellon University, Pittsburgh, PA, USA; Center for the Neural Basis of Cognition, Carnegie Mellon University & University of Pittsburgh, Pittsburgh, PA, USA; Center for Neuroscience, University of Pittsburgh, Pittsburgh, PA, USA; Machine Learning Department, Carnegie Mellon University, Pittsburgh, PA, USA; Department of Anatomy and Neurobiology, Boston University, Boston, MA, USA; Department of Biomedical Engineering, Carnegie Mellon University, Pittsburgh, PA, USA; Department of Electrical and Computer Engineering, Carnegie Mellon University, Pittsburgh, PA, USA; Department of Bioengineering, University of Pittsburgh, Pittsburgh, PA, USA

**Keywords:** Reward, visual cortex, neuronal population coding, attention

## Abstract

Even professional athletes sometimes fail to perform when it matters most. This phenomenon, known as choking under pressure, has been observed in a variety of motor and cognitive tasks. Can inordinately large rewards influence even sensory processing, and if so, what might be its neural basis? To probe this, we studied rhesus monkeys performing a challenging color change detection task and recorded from populations of neurons in visual area V4. We found that monkeys underperformed when offered rare “jackpot” rewards and uncovered associated detrimental alterations in V4 population activity. These findings were consistent with misallocated spatial attention, providing a potential neural basis for perceptual choking under pressure.

## INTRODUCTION

Increasing incentives tends to improve performance^1–4^. Colloquially, this is the difference between practice versus game time: when the potential reward is high, performance should be optimal. However, increasing the stakes too much can lead to a paradoxical decrease in performance, commonly called “choking under pressure”^5–7^. Previous studies have documented choking under pressure in humans performing a variety of motor^8–15^ and cognitive^16–20^ tasks. A more recent body of work has begun to examine the neural underpinnings of choking under pressure; human neuroimaging studies have found neural correlates of choking under pressure in a variety of brain areas^21–26^ including those canonically linked to reward processing.

Monkeys, like humans, can choke under pressure^27^. When offered a rare, uncharacteristically large “jackpot” reward, monkeys’ motor performance significantly declined. This drop in performance in an arm movement task correlated with weaker preparatory signals in motor cortex activity preceding movement, pointing to one potential neural basis of choking under pressure^28^. However, given the wide array of behavioral processes that exhibit choking under pressure, it’s unlikely that motor preparatory signals are the only ones corrupted by extraordinary rewards. Reward modulates many brain areas involved in processes such as executive function^29–35^ and sensory perception^36–40^. Might any of these processes contribute to choking under pressure? If so, even the most elemental perceptual processes, such as visual detection, might be adversely affected by inordinately high rewards.

We sought to uncover whether perceptual systems contributed to choking under pressure by probing visual behavior and its neural correlates in mid-level visual cortical area V4. To do this, we developed a task combining a challenging perceptual change detection (a color change) and a precise arm movement to a target, which also contained the pre-cued reward at stake for each trial. In addition to previously established motor performance deficits, monkeys’ detection performance for jackpot rewards was significantly worse than for lesser rewards, suggesting that errors in sensory processing also contributed to choking under pressure. We then examined neural correlates that might underlie this behavior and found that visual stimulus encoding in V4 was weaker on jackpot trials. In contrast, V4 exhibited stronger encoding of the location of the reward cue on jackpot trials, even though the color change occurred far from that location. These findings support the hypothesis that the animals were distracted from the stimulus and oriented toward the reward cue on jackpot trials; or, they had their “eyes on the prize”. Our results thus highlight a distinct perceptual component of choking under pressure originating from the misallocation of attentional resources.

## RESULTS

We trained three monkeys to perform a challenging task in which they had to detect a subtle color change of a contrast-modulated sinusoidal grating (illustrated as gray to blue; Figure 1a) and then report this detection by quickly moving a joystick-controlled cursor (green circle) to a small movement target (white circle). With each contrast cycle of the grating (after a minimum stimulus presentation period), there was an equal chance of a color change occurring, encouraging the animals to focus on the grating and wait to detect the color change. On each trial, the amount of reward available upon success was indicated to the animal using pre-learned cues (isoluminant letters collocated with the movement target). On 95% of trials, a small (S), medium (M), or large (L) liquid reward was available. On a rare 5% of trials, an exceptionally-sized “jackpot” reward (J) was available (Figure 1b). To elicit choking under pressure, both the perceptual and motor components of the task were titrated to be difficult (Methods, Figure S1). The magnitude of the color change was varied randomly from trial-to-trial over a fixed range (4 possible color change magnitudes; Figure 1c), and the movement target diameter and movement time limit were tightly constrained. We first asked whether the errors associated with choking under pressure extended to the perceptual domain as well as the motor domain.

**Figure 1:**
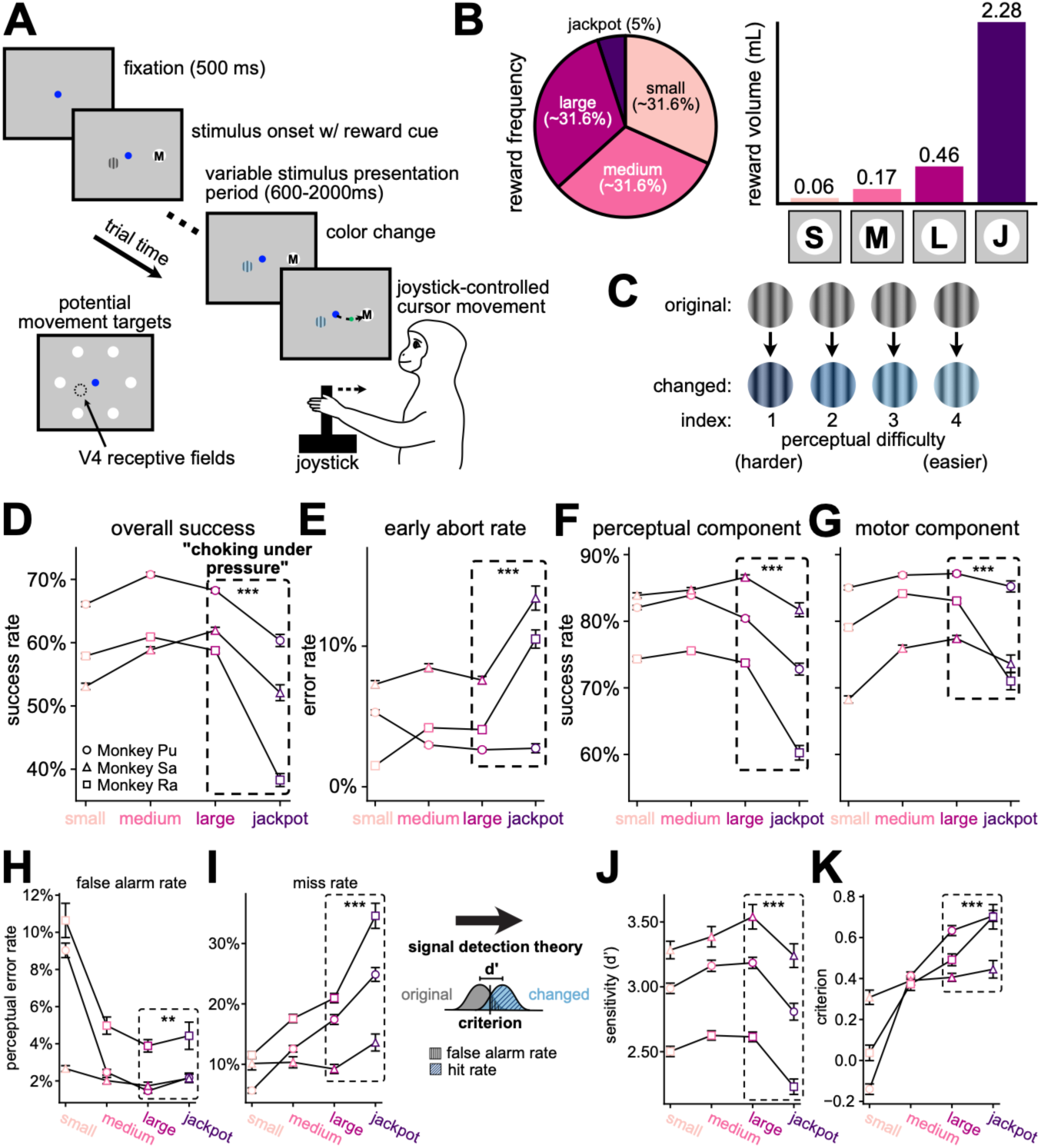
Distinct perceptual and motor components comprised choking under pressure. (A) Animals performed a color change detection task for variable, pre-cued rewards (Methods). (B) Left, reward was offered at equal frequency for small, medium, and large (∼31.6% of trials) and rarely for jackpot (5%). Right, reward cues and corresponding volumes from an example monkey (Monkey Pu) are shown in milliliters (Table S1). (C) Illustrative example of color change magnitudes. The original hue (top) was always the same, and the changed hue (bottom) was one of four graded difficulties on each trial, indexed from 1 (harder) to 4 (easier). (D) Overall task performance (rewarded trials / total trials) as a function of reward. Each point indicates the performance +/-standard error (SE) across trials in each condition, computed assuming Bernoulli-distributed success rates (note that error bars are sometimes smaller than the plotted point). Monkeys choked under pressure (large success rate > jackpot, p<0.0001, pooled across animals; Monkey Pu: p<0.0001, circles; Monkey Sa: p<0.0001, triangles; Monkey Ra: p<0.0001, squares; binomial test). (E) Early abort rates (aborted trials / total trials) increased on jackpot trials compared to large reward trials (p<0.0001, pooled across animals; Monkey Pu: p=0.38, Monkey Sa: p<0.0001, Monkey Ra: p<0.0001; binomial test). (F) Detection performance (hits / non-aborted trials; Methods) as a function of reward. Monkeys choked under pressure on the perceptual component of the task (p<0.0001, pooled across animals; Monkey Pu: p<0.0001, Monkey Sa: p<0.0001, Monkey Ra: p<0.0001; binomial test) on all color change magnitudes (Figure S2). Conventions as in D. (G) Motor performance (rewarded trials / hits) as a function of reward. Monkeys choked under pressure on the motor component of the task (p<0.0001, pooled across animals; Monkey Pu: p=0.0084, Monkey Sa: p=0.0017, Monkey Ra: p<0.0001; binomial test). Conventions as in D. (H) False alarm rates increased on jackpot trials compared to large reward trials (p=0.0066, pooled across animals, n=93 sessions; Monkey Pu: p=0.00088, n=42, circles; Monkey Sa: p=0.051, n=25, triangles; Monkey Ra: p=0.23, n=26, squares; paired t-test). Each point indicates the mean +/-SE across sessions in each animal. (I) Miss rates increased on jackpot trials compared to large reward trials (p<0.0001, pooled across animals, n=93 sessions; Monkey Pu: p<0.0001, n=42; Monkey Sa: p=0.00038, n=25; Monkey Ra: p<0.0001, n=26; paired t-test). Conventions as in H. (J) Perceptual sensitivity (d’) decreased on jackpot trials compared to large reward trials (p<0.0001, pooled across animals, n=93 sessions; Monkey Pu: p<0.0001, n=42; Monkey Sa: p=0.0020, n=25; Monkey Ra: p<0.0001, n=26; paired t-test). Conventions as in H. (K) Criterion increased on jackpot trials compared to large reward trials (p<0.0001, pooled across animals, n=93 sessions; Monkey Pu: p=0.0052, n=42; Monkey Sa: p=0.18, n=25; Monkey Ra: p=0.00022, n=26; paired t-test). Conventions as in H. ***p < 0.001 by binomial test (D-G) or paired t-test (I-K) on trials/sessions pooled across animals. ** p < 0.01 by paired t-test on sessions pooled across animals (H).

### Distinct perceptual and motor components comprised choking under pressure

As in previous work^27^, animals exhibited a marked decrease in overall success rate on our sensorimotor task when offered a jackpot reward, compared to their performance on large reward trials (Figure 1d). To determine why they failed, we subdivided their errors into different categories. First, animals were required to maintain fixation until the color change, which always occurred during or after the third contrast cycle (≥600ms from stimulus onset). However, when jackpot rewards were presented, we observed a significant increase in trials in which the animal aborted the trial by looking away from the fixation point, and typically to the reward cue, early in the stimulus presentation period (<600ms, “early aborts”, Figure 1e). If animals maintained fixation through this initial period, they had the opportunity to detect the color change. We found that performance on this perceptual component of the task significantly decreased on jackpot trials compared to large reward trials (Figure 1f). If the animals correctly detected the color change, they were required to report their detection by quickly moving to the movement target. Given successful detection, we observed a further decrease in motor performance on jackpot trials (Figure 1g), consistent with our previous work^27^.

Failures on the perceptual component of the task could be attributed to a variety of sources, including errant decision-making, motor preparation, or true perceptual failures. To determine whether these failures were truly related to perception, we subdivided them into false alarms (Figure 1h) and misses (Figure 1i). We then applied signal detection theory to assess changes to their perceptual sensitivity (d’) and criterion (Methods). If perception contributed to choking under pressure, we would expect to see a decrease in sensitivity. This is indeed what we found: sensitivity was significantly worse on jackpot trials compared to large reward trials (Figure 1j). Interestingly, criterion increased monotonically with reward size (Figure 1k), indicating that animals adopted a strategy in which they were more cautious when larger rewards were offered.

To further isolate the perceptual component of choking under pressure, we trained two animals to perform another variant of the task in which the motor effector was changed to be a comparatively simple visually-guided saccade to the movement target. Even without challenging motor demands, the impact of perceptual choking under pressure remained (Figure S3). Taken together, these behavioral results reveal that choking under pressure extends into the perceptual domain.

### Jackpot rewards increased correlated variability in the V4 population response

We next investigated a potential neural basis for the perceptual failures in the responses of populations of neurons in visual area V4 (Figure 2a, all neural data from Monkey Pu and Monkey Ra). If V4 reflected the perceptual component of choking under pressure, we might expect to see alterations to the sensory information about the stimulus provided by V4 on jackpot trials. This might occur through a decrease in response gain, leading to a reduction in stimulus-evoked firing rates^41^. We thus investigated how reward changed V4 firing rates during the stimulus presentation period, just before a color change might occur. While large rewards produced reliable increases in stimulus-evoked firing rates compared to small rewards (Figure 2b-c), jackpots produced inconsistent changes in population average firing rates across animals (Figure 2d). Even so, within each animal, reward information was decodable in V4 throughout the stimulus presentation period (Figure S4), indicating that jackpot rewards meaningfully altered the V4 population response, but not in a manner consistent with reduced stimulus response gain. This led us to consider other ways in which jackpot rewards might impact population responses in V4.

**Figure 2:**
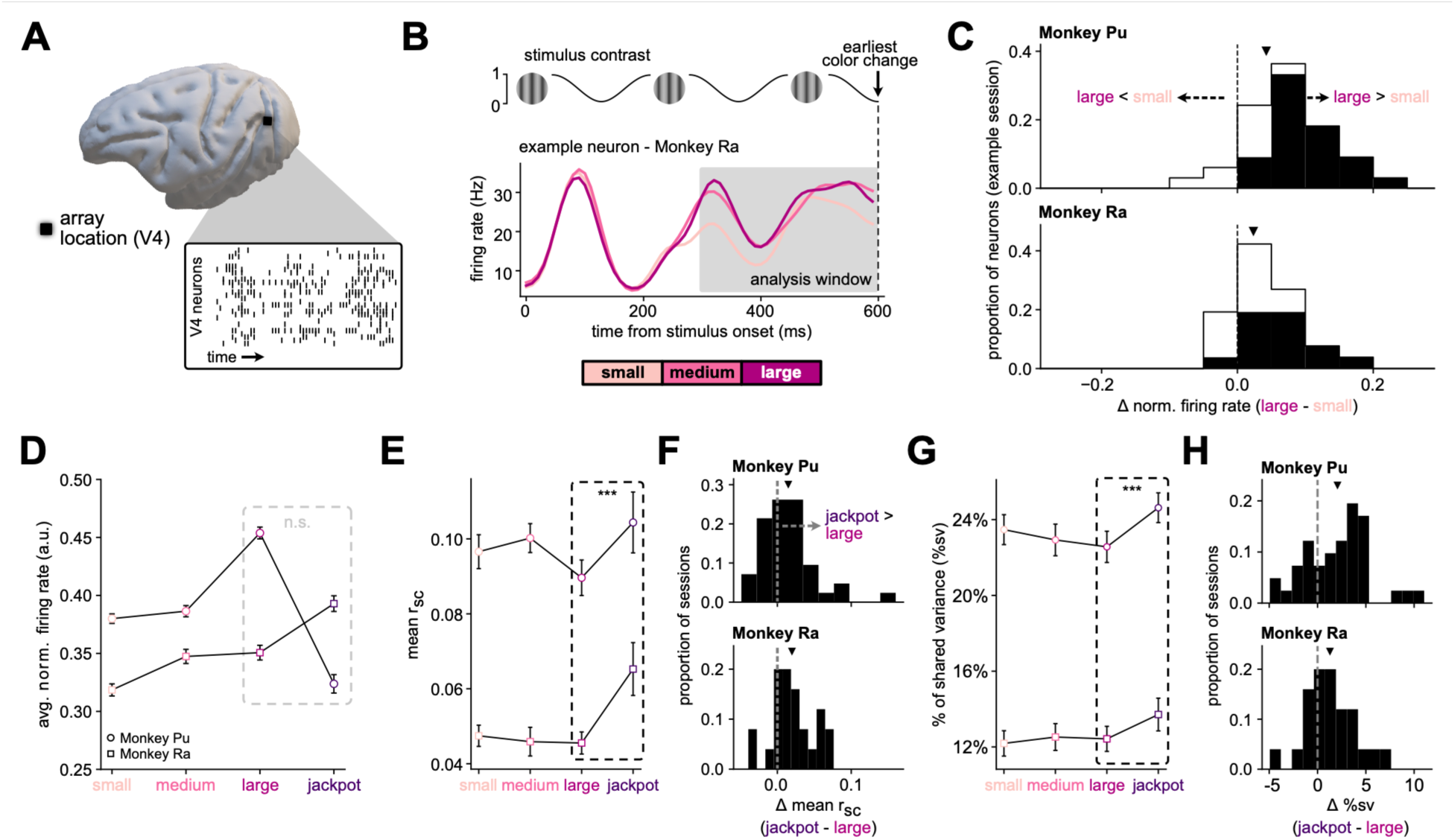
Jackpot rewards increased correlated variability in the V4 population response. (A) Example raster from a multi-electrode array implanted in V4 in Monkey Ra. (B) Top, time-course of stimulus contrast (Methods). Bottom, trial-averaged activity split by reward condition, smoothed with a Gaussian kernel (width=15ms), for a typical neuron in an example session from Monkey Ra. Analysis window for C-D shown in the shaded area. (C) Per-neuron difference (Δ) in normalized firing rate between small and large reward trials in an example session (Monkey Pu: top, n=33 neurons; Monkey Ra: bottom, n=26 neurons). Proportion of neurons that significantly increased or decreased their firing rate are filled with solid bars (Monkey Pu: 72.7% neurons showed significantly increased firing rate for large reward trials compared to small reward trials, Monkey Ra: 50.0%; Mann-Whitney U test). Mean of each distribution marked with a triangle. (D) Population average normalized firing rate during the stimulus presentation period (300-600ms after stimulus onset). Each point indicates the mean +/-SE across sessions in each animal. The mean rate did not consistently increase from large to jackpot trials (p=1.0, pooled across animals, n=67 sessions; Monkey Pu: p=1.0, n=42, circles; Monkey Ra: p<0.0001, n=25, squares; paired one-tailed t-test). (E) The mean *r_sc_* across pairs of V4 neurons increased from large to jackpot trials (p<0.0001, pooled across animals, n=67 sessions; Monkey Pu: p=0.0068, n=42, circles; Monkey Ra: p=0.0014, n=25, squares; paired t-test). Each point indicates the mean +/-SE across sessions in each animal. (F) Histograms indicating the individual session differences (Δ) in mean *r_sc_* from large to jackpot trials. Mean of each distribution marked with a triangle (Monkey Pu: mean Δ = 0.015, top; Monkey Ra: mean Δ = 0.020, bottom). Each histogram shows the individual session values of the data shown in E. (G-H) %sv from an FA model increased from large to jackpot trials (p<0.0001, pooled across animals, n=66 sessions; Monkey Pu: mean Δ = 2.06%, p=0.00020, n=41; Monkey Ra: mean Δ = 1.29%, p=0.0094, n=25; paired t-test). Conventions as in E-F. ***p < 0.001 by paired t-test pooled across animals (E and G).

Firing rate shifts are not the only way that alterations in neural activity may be linked with behavior: reward might also impact the patterns of correlated variability among neurons. Trial-to-trial correlated variability has been linked to the fidelity of sensory coding^42–44^, and decreases in this variability are associated with performance improvements in various task contexts such as learning^45–47^, attention^48–53^, and experience^54^. If perceptual choking under pressure was a result of reduced sensory information, we might also expect to see an increase in correlated variability among neurons. To investigate this, we considered both correlated variability among V4 neuron pairs (mean *r_sc_*, Methods; Figure 2e-f) and a population-level covariability metric (percent shared variance, %sv, Methods; Figure 2g-h) during the stimulus presentation period just before a potential color change. In both cases, we observed significant increases in correlated variability on jackpot trials relative to large reward trials. Thus, even before the color change occurred, we observed a modulation of the V4 population response that could impact the fidelity of stimulus coding by reducing the signal-to-noise ratio^55^.

### Jackpots reduced color change signals in V4

We next analyzed V4 population activity during a time period critical to perceptual success: the time of the color change. Due to the limited number of the rare jackpot trials in each experimental session, we combined trials across sessions using a neural “stitching” algorithm^56^ (Methods). To isolate neural activity that corresponded to the evidence that a color change occurred, we defined a “signal axis” using linear discriminant analysis that best separated V4 activity when the color had changed (Signal Present; Figure 3a, blue) from when it had not changed (Signal Absent; gray). Along the signal axis, activity in the Signal Present trials increased after the color change (Figure 3b, blue) relative to the Signal Absent trials during the same period (gray).

**Figure 3:**
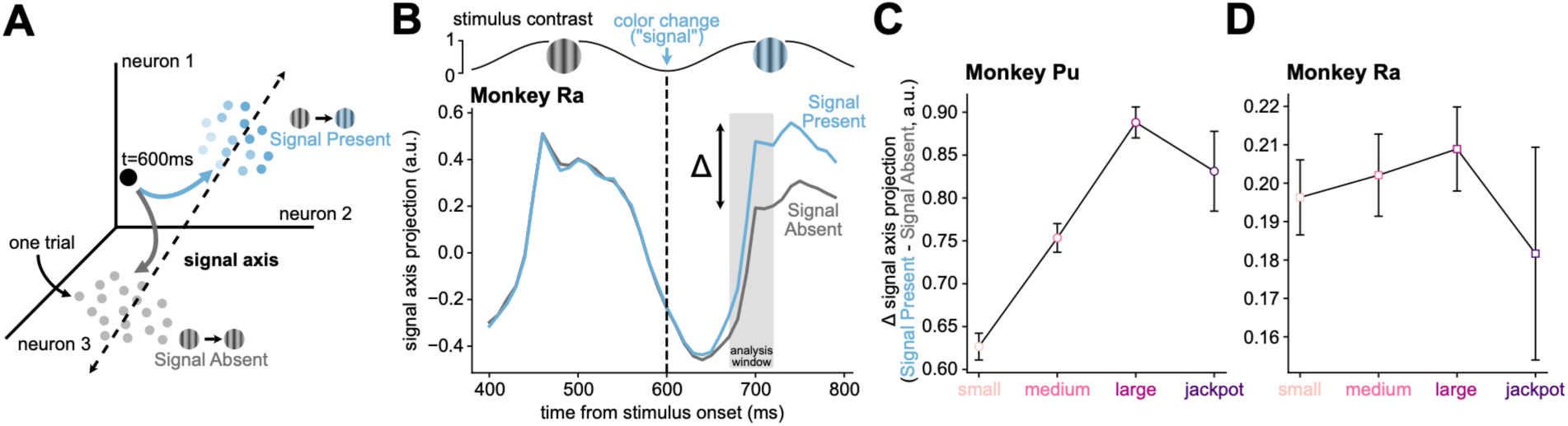
Jackpots reduced color change signals in V4. (A) Schematic defining a “signal axis” that best separated neural activity when the color changed (Signal Present, blue points) from when it had not (Signal Absent, gray points). Each coordinate axis represents the activity of one neuron. (B) Projections along the signal axis (Monkey Ra) for Signal Present (blue) and Absent trials (gray) diverged shortly after the change occurred (600ms, dashed line; also indicated by grating schematic at top). Traces indicate mean +/-SE across trials. Analysis window for C shaded in gray. (C-D) The difference (Δ) between signal axis projections from Signal Present to Absent trials was lower for jackpot rewards than for large rewards (p=0.041, pooled across animals, n=56 sessions; C: Monkey Pu, p=0.081, n=33; D: Monkey Ra, p=0.20, n=23). Each point indicates the difference +/-SE across trials.

To quantify the color change signal in V4, we computed the difference (Δ) in signal axis projections between Signal Present and Absent trials in each reward condition (Figure S5). By assessing this difference, we isolated the impact of reward on the color change signal above and beyond any effects that were present before the color change (Signal Absent). If jackpot rewards reduced available sensory information compared to large rewards, we would expect to see a decrease in the color change signal along this axis.

We computed the average Δ separately for each reward condition and found a reduction on jackpot trials compared to large reward trials (Figure 3c-d). In short, the color change signal in V4 was smaller on jackpot trials than on large reward trials.

### Jackpots drew spatial attention away from the stimulus to the movement target

Both the drop in color change signal (Figure 3) and the increase in correlated variability (Figure 2) have negative implications for downstream readouts of V4: reducing the available sensory information before the animal could report the color change detection. Previous studies of visual cortex have shown that directing spatial attention away from a stimulus in the receptive fields results in increased correlated variability^48,49,51–53^. Thus, one explanation of our results is that spatial attention was not properly allocated to the stimulus on jackpot trials and that the animal was distracted. We sought to determine whether, on jackpot trials, the animals were less able to keep their attention on the stimulus because they were distracted by the movement target containing the high-value jackpot cue.

During the stimulus presentation period, another object was present on the screen: the movement target containing the reward cue (Figure 1a). If the animal properly allocated attention to the stimulus (stimulus was “attended”; Figure 4a, top), the impact of the movement target on the visual response would be low, as the movement targets were located well outside of the classical receptive fields of the recorded neurons (Methods). However, shifts in attention are known to alter responses to stimuli in the extra-classical surround of V4 neurons^57^. Thus, if the animal inappropriately attended to the movement target instead of the stimulus (stimulus was “unattended”; bottom), the impact of the movement target on the visual response would be high.

**Figure 4:**
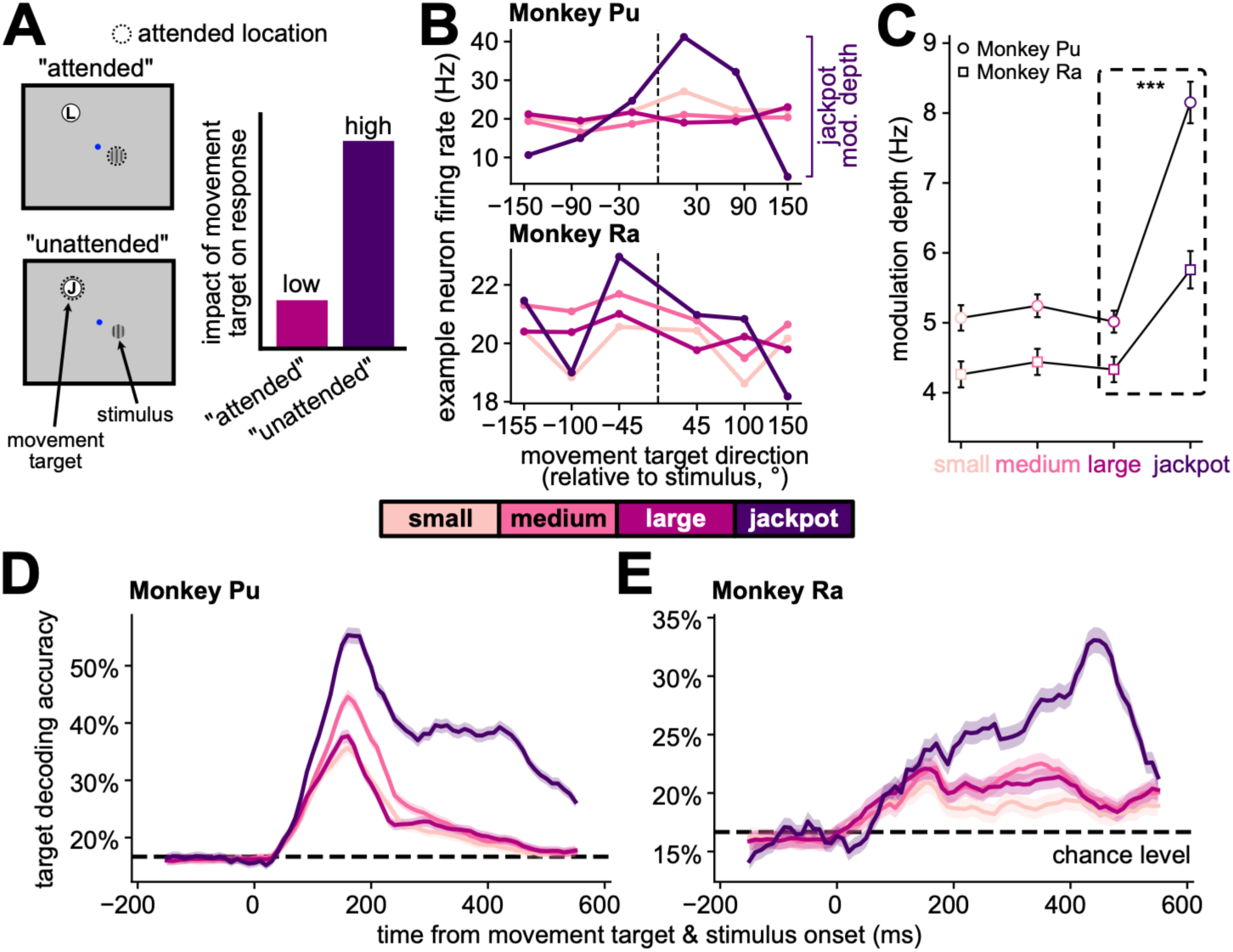
Jackpots drew spatial attention away from the stimulus to the movement target. (A) Left, two hypothesized loci of spatial attention (dashed circles). Left top, stimulus is “attended”. Left bottom, stimulus is “unattended” when attention is directed to the movement target. Right, impact of the movement target on the visual response should be low when attention is on the stimulus and high when it is not. (B) Firing rates of example neurons to each movement target location under different reward conditions (Monkey Pu: top, Monkey Ra: bottom). The horizontal axis position indicates the relative angle between the direction of the movement target and the direction of the stimulus, in degrees. Modulation depth is the difference between the maximum and minimum average firing rates to the different movement target locations. (C) Modulation depth was highest on jackpot trials (p<0.0001, pooled across animals, n=52 sessions; Monkey Pu: p<0.0001, n=28, circles; Monkey Ra: p<0.0001, n=24, squares; paired t-test). Each point indicates the mean +/-SE across sessions in each animal. ***p < 0.001 by paired t-test pooled across animals. (D-E) The movement target location was decodable from V4 population activity, and highest on jackpot trials (D: Monkey Pu, E: Monkey Ra). Shaded area indicates the SE across folds and repeated subsamples (Methods). Chance level is ∼16.6% (6 movement target locations, presented equally). The horizontal axis position indicates the center of the 100ms bin used to train the decoder.

To test this hypothesis, we assessed how strongly the movement targets impacted stimulus-evoked firing rates for each V4 neuron by computing their modulation depth – that is, the change in firing rate to the different movement target locations during the stimulus presentation period (Figure 4b). If the modulation depth were large, it would indicate increased attention to the movement target as opposed to the stimulus. Aggregated across sessions, we found that modulation depth was highest on jackpot trials (Figure 4c). This indicates that on jackpot trials, movement targets had a higher impact on V4 firing rates during the stimulus presentation period (a time before the movement target was behaviorally relevant), supporting the hypothesis that attentional resources may have been drawn toward the movement target.

Next, we studied the impact of the movement target on the response of the entire population of V4 neurons. We trained a linear decoder to predict the location of the movement target from stitched V4 population activity during the stimulus presentation period. If attention were directed to the stimulus, we would expect a minimal impact of the movement target on the V4 population response and thus poor decoding of the movement target location. On small, medium, and large reward trials, cross-validated decoding performance fell to near chance levels (∼16.6% for 6 movement target locations) early in the stimulus presentation period (t<300ms; Figure 4d-e), indicating that the movement target was not substantially impacting V4 responses. In contrast, on jackpot trials decoding performance was sustained throughout the stimulus presentation period and was higher than for all other rewards. Together, these results support the hypothesis that spatial attention resources were drawn to the movement target, altering V4 population activity in a manner consistent with the distinct perceptual component of choking under pressure we observed.

## DISCUSSION

In this study, we report that monkeys, like humans, choked under pressure in multiple ways, most notably in terms of worse perceptual performance: when given a challenging color change detection, performance for jackpot rewards was significantly worse than for lesser rewards. We examined the neural underpinnings of this perceptual choking under pressure by focusing on changes in visual area V4. We found multiple lines of evidence that supported the hypothesis that animals were distracted away from the stimulus to the movement target on jackpot trials (i.e., animals had their “eyes on the prize”). First, there were consistent increases in trial-to-trial variability. Second, color change signals were reduced. Lastly, the movement target containing the reward cue had the greatest impact on V4 activity on jackpot trials compared to other rewards. We thus propose a theory for perceptual choking under pressure in which reward modulates the allocation of attention, shaping V4 activity and, in turn, behavior.

Reward has long been linked to attention. Rewarding stimuli can capture exogenous attention^58^, a phenomenon known as value-driven attentional capture^59^. In addition, they also influence the allocation of endogenous attention, where more rewarding stimuli receive more attentional resources^60^. Indeed, researchers often vary reward contingencies to manipulate endogenous spatial and feature attention in experiments^61^. In our task, the appearance of jackpot rewards was associated with many of these aspects of attentional modulation. The increase in early aborts on jackpot trials can be interpreted as value-driven attentional capture that was unable to be suppressed, or a breakdown in response inhibition. A breakdown in response inhibition could also explain our movement target decoding results, which support the argument that the highly rewarding jackpot resulted in attention being deployed to the wrong object (the movement target, as opposed to the stimulus). Misallocated spatial attention is also consistent with the decreased perceptual sensitivity and increased criterion revealed by our signal detection theory analysis. The animals were worse at detecting color changes on the now “unattended” stimulus and responded less frequently or less impulsively, a finding recapitulated in recent modeling work^62^. These findings are also consistent with a shift in the spatial locus of attention (due to multiple competing stimuli) or a reduction in overall attentional intensity^63^. Our results support the hypothesis that jackpot rewards led to inappropriate allocation of attentional resources.

Attention has been shown to impact multiple aspects of visual cortical activity (including V4) and, in turn, behavior. Endogenous spatial or feature attention tends to increase firing rates to the attended location or feature^64–66^, and filters out responses to non-attended stimuli^67–69^. Changes in receptive field (RF) size as well as interactions with the region surrounding the RF (the visual surround)^57,67,70–72^ are also known to occur with changes in spatial attention – attention to a point within the RF tends to shrink the size of the RF around that point, and attention to a region outside the RF tends to expand its size. While our study was not designed to probe RF properties explicitly, the changes in movement target modulation depth we observed are indicative of changes in RF size or interactions with the surround. Beyond single-neuron properties, decreases in pairwise and population-level correlated variability are associated with the deployment of spatial attention resources^48,49,73^, consistent with what we observed. Reduced variability has been proposed to improve the fidelity of sensory coding in neuronal populations^42,74^, and such modulations of V4 have been linked to changes in behavioral performance^48,49,51,52^. Other behaviorally relevant modulations of V4 activity have been shown in contexts such as learning tasks^45^, experience with recent visual stimuli^54^, and changes in task difficulty^63,75–77^. In these studies, V4 activity is typically modulated in a manner that corresponds with improved behavioral performance. In our work, however, we observed a modulation of V4 that decreased performance, because attentional resources were drawn away from the relevant stimulus.

To what extent is impaired attention an explanation of choking under pressure in other paradigms? Distraction, or a lapse in task-relevant attention, is a prominent psychological theory for choking under pressure^15,16,78^, and the reduced perceptual performance we observed is consistent with the animals being distracted. Almost every task requires perception, and thus attention’s influence on perceptual capabilities could impact performance on almost any task. In perceptual tasks, however, the impact of attention is greatest when the perceptual component is difficult, and minimal when easy^76,79–81^. In a task with trivial perceptual demands (like our previous study of motor cortex^28^ and many other studies of choking under pressure), we might not expect spatial attention to play as dominant a role as in the present work. However, attention includes processes that are much broader than spatial allocation of perceptual resources across the visual field. It may be that attention’s contributions to choking under pressure occur through not only perceptual sensitivity changes^48,82^ but also through higher-order cognitive processes such as decision-making^36,67,83,84^, multi-sensory integration^85^, or the suppression of distractors^86,87^. For example, impairment of the attentional control of a basketball player might make them susceptible to distraction by a booing crowd when attempting a free throw, though their visual perception of the hoop remains unaffected.

While attention may be obligatorily yoked with changes in reward or stakes^61^, the effects of reward are not limited to modulation of attention. Reward signals spread throughout the brain and impact a wide array of neural processes^88^ including sensory perception^36–40^, executive function^29–35^, and motor planning^28,89,90^. Any of these systems might be subject to dysregulation when extraordinary rewards are at stake. Consistent with this view, human neuroimaging studies have found correlates of choking under pressure throughout the brain^21–26^. A complete account of choking under pressure must therefore incorporate an understanding of the impacts of reward on brain-wide computations.

## ACKNOWLEDGMENTS

The authors would like to thank Samantha Nelson and Emilio Salazar-Gatzimas for help with animal training, Emily Oby for jointly performing array implant surgeries, our animal care staff for their dedication in caring for the animals used in this study, and Emilio Salazar-Gatzimas and Patricia Stan for helpful discussions.

M.E.M. was supported by NIH T32 EB029365. A.N.C. was supported by the Mellon Predoctoral Fellowship. C.K. was supported by NIH T32 NS126122. A.L.S. was supported by NIH F32 NS144018 and the Boston University Center for Systems Neuroscience Postdoctoral Fellowship. H.M. was supported by JASSO G2599999939N. B.M.Y., A.P.B., M.A.S., and S.M.C. were supported by NSF NCS DRL 2124066/2123911. B.M.Y., A.P.B., and S.M.C. were supported by NIH R01 NS105318, and NIH R01 NS129098.

## AUTHOR CONTRIBUTIONS

Conceptualization, M.E.M., A.N.C., C.K., A.L.S., H.M., B.M.Y., A.P.B., S.M.C., and M.A.S.; Data curation: M.E.M., A.N.C., and C.K.; Formal analysis, M.E.M.; Funding acquisition, B.M.Y., A.P.B., S.M.C., and M.A.S.; Investigation, M.E.M., A.N.C., and C.K.; Methodology, M.E.M., A.N.C., C.K., A.L.S., H.M., B.M.Y., A.P.B., S.M.C., and M.A.S.; Resources, B.M.Y., A.P.B., S.M.C., and M.A.S.; Supervision, B.M.Y., A.P.B., S.M.C., and M.A.S.; Validation, M.E.M.; Visualization, M.E.M.; Writing – original draft, M.E.M.; Writing – review & editing, M.E.M., A.N.C., C.K., A.L.S., H.M., B.M.Y., A.P.B., S.M.C., and M.A.S.;

## SUPPLEMENTAL INFORMATION

Document S1. Supplemental Figures S1–S9 and Supplemental Tables S1–S2

## METHODS

### Experimental Model

Three male rhesus macaque monkeys (*Macaca mulatta*) were used in this study. For the Hand Task, they were 8 (Monkey Pu) and 11 (Monkey Ra and Monkey Sa) years old at the time of data collection. For the Eye Task, they were 10 (Monkey Pu and Monkey Ra) years old at the time of data collection. The animals were housed singly in a room operating on a standard 12-hour light/dark cycle and were provided with an enhanced enrichment program. All experimental procedures were approved by the Institutional Animal Care and Use Committee of Carnegie Mellon University and were in accordance with the United States National Research Council’s Guide for the Care and Use of Laboratory Animals.

### Surgical preparation

Each animal was surgically implanted with a titanium headpost, which was affixed to the skull using titanium screws. This was necessary to limit head movement during recording sessions such that the position of the eyes could be tracked and neural activity recorded. In a separate procedure, a subset of animals (Monkey Pu and Monkey Ra) were implanted with a 96-electrode “Utah” array (Blackrock Neurotech, Salt Lake City, UT). The electrode arrays were placed over visual area V4, left hemisphere for Monkey Pu and right hemisphere for Monkey Ra. Monkey Sa did not have an implant in V4, and only behavioral data from this animal is included. Surgeries were performed in sterile conditions under general anesthesia using isoflurane.

### Behavioral Tasks

All visual stimuli were shown on a luminance-corrected cathode ray tube monitor with 1024×768-pixel resolution and a refresh rate of 100 Hz, positioned 46 cm (Monkey Pu and Monkey Sa) or 57 cm (Monkey Ra) away from the animal’s eyes. Stimuli were generated with custom-designed software in MATLAB (Mathworks, Natick, MA) that utilized the Psychophysics toolbox extensions^92,93^.

### Receptive field mapping

We mapped receptive fields (RFs) of the V4 neurons to determine the appropriate size and location for the stimulus used in the change detection task. Sinusoidal gratings of four possible orientations ([0°, 45°, 90°, 135°] from horizontal) were displayed one at a time in a grid of positions that covered the likely RF area based on the anatomical location of the implant (lower contralateral quadrant of the visual field). We identified the center and extent of each neuron’s RF by analyzing the mean response to gratings presented at each location. The size and position of the final stimulus for the change detection task were chosen to cover the aggregate of the V4 RFs recorded by the array. For the Hand Task, the stimulus center location (X, Y) in degrees of visual angle (d.v.a.) was (+2.0, -1.1) for Monkey Pu and (-1.1, -1.8) for Monkey Ra. For the Eye Task, the stimulus center locations were (+2.8, -1.1) for Monkey Pu and (+5.5, -10.9) for Monkey Ra. Negative X coordinates indicate that the stimulus was centered to the left of fixation, and negative Y coordinates indicate that the stimulus was centered below fixation. For the Hand Task, the stimulus diameter in d.v.a. was 2.2 for Monkey Pu and 1.8 for Monkey Ra. For the Eye Task, the stimulus diameter was 2.0 for Monkey Pu and 5.8 for Monkey Ra. Monkey Sa did not have a V4 implant (behavior only) so no receptive field mapping or stimulus parameter titration was performed.

### Change detection task (Hand Task)

Each animal was trained to perform a challenging variant of a visual change detection task for variable rewards (Figure 1a). Briefly, a visual stimulus and a movement target containing a learned reward cue appeared on the screen. After a variable length stimulus presentation period, the color of the stimulus changed, serving as the go-cue for the animal to move to the movement target. Upon success, they received the cued reward (small, medium, large, or jackpot). In total, we recorded 42 sessions from Monkey Pu, 25 sessions from Monkey Sa, and 26 sessions from Monkey Ra. The details of the task were as follows.

#### Stimulus and movement target presentation

Following a 500ms fixation period at the center of the screen, the stimulus for the color change detection (an oriented sinusoidal grating) appeared in the animal’s V4 RF. At the same time, the movement target for the report appeared at one of 6 locations in the periphery (locations were spaced to not be in the same direction as the stimulus, and were more eccentric than the stimulus). The movement target indicated the location at which the animal would complete their report of the color change and contained a visual cue as to the level of reward the animal would receive upon success. The reward cues were isoluminant letters which indicated the size of the reward (S: small, M: medium, L: large, J: jackpot). The letters were drawn as filled pixels in a 9×9 grid that covered the area of the movement target, with each letter filling the same number of pixels (and thus the average luminance across all the letters was the same). On each trial, there was a 5% chance of a jackpot reward being presented. If a jackpot was not presented, a small, medium, or large reward was equally likely to appear (∼31.6%). For reward volumes, see Table S1.

#### Perceptual component

A variable length stimulus presentation period followed, during which the grating was contrast-modulated. The grating was first presented at full contrast, and one contrast cycle was a full period of a sinusoidal function in which the grating faded to zero contrast and returned to full contrast (240ms). During or after the third contrast cycle, the stimulus could change color. Color changes only occurred at times when the grating contrast was zero (indistinguishable from monitor background). Thus, color changes could occur at 600ms from stimulus onset (halfway through the third contrast cycle), and every 240ms thereafter. The distribution over trials of times at which the color changed followed a discretized exponential distribution, where at every contrast cycle after the minimum, the grating had a 35% chance to change color. This approximated a flat hazard rate across possible color change times, such that the color change was equally likely to occur at every contrast cycle given that it had not yet occurred. A maximum stimulus presentation period length was set at 2500ms. The magnitude of the color change could take one of four values on each trial. For color change magnitude values, see Table S2. For detection performance as a function of change magnitude index, see Figure S1.

#### Motor component

Two variants of the task were run: one in which the movement to the target was performed using a joystick-controlled cursor (hand movement, termed “Hand Task”), and the other in which the movement to the target was a saccadic eye movement (termed “Eye Task”). In each case, the motor requirements were as follows. When the animal detected that the color had changed, they initiated their report by quickly moving the effector (cursor or eyes) to the movement target. A movement from the central fixation point needed to occur within a tight window (given the perceptual difficulty) to be considered a perceptual hit, otherwise it was considered a miss. For the Hand Task, the animals were required to react between 180-480ms (Monkey Pu), 80-450ms (Monkey Sa), or 190-550ms (Monkey Ra). For the Eye Task, the animals were required to react within 100-500ms (Monkey Pu) or 100-450ms (Monkey Ra). After the animal reacted, the effector was required to reach the movement target within a limited time. For the Hand Task, the movement time limit was 400ms (Monkey Pu), 230ms (Monkey Sa), or 325ms (Monkey Ra). For the Eye Task, the movement time limit was 100ms (Monkey Pu) or 60ms (Monkey Ra). The time limits for the motor component were titrated for each animal after initial training on the task, which was less constrained. Note, the movement time limits for the Eye Task were faster than those for the Hand Task, as the eyes move much faster than the hands. Upon acquiring the movement target, the effector needed to be held at the movement target for 250ms, after which the cued reward was delivered.

#### Variant for Monkey Sa

For Monkey Sa, who did not have a V4 implant (behavior only), the central fixation point served as the stimulus for color change detection. No grating appeared on the screen. Movement targets appeared in one of 8 evenly spaced locations in the periphery. Following a variable length stimulus presentation period, the central fixation point changed color, which served as the animal’s go-cue to report to the movement target. Color change magnitudes were one of six graded difficulties (instead of four). Stimulus presentation period lengths were exponentially distributed with a mean of 325ms, after a minimum of 500ms. All other task mechanics were as described above.

#### Other task manipulations

There were additional trial-by-trial manipulations beyond the movement target location, reward, and color change magnitude described above. However, they were not relevant for the analyses presented in this work, and so we combined all trial types. We describe the additional manipulations here for completeness. On 50% of trials, the movement target diameter was increased by 1.5x. On a longer timescale, perceptual difficulty was shifted up or down in blocks of 50-200 trials (depending on the animal). For Monkey Pu and Monkey Ra, during easy perceptual blocks, color change magnitudes were presented with frequencies of 10%, 20%, 30%, and 40% for color change magnitude index 1, 2, 3, and 4, respectively. During hard perceptual blocks, these frequencies were reversed such that color change magnitude 1 (hardest) was presented with 40% frequency. For Monkey Sa, each color change magnitude was equally likely to appear on a given trial, subject to the following block-wise difficulty constraint: the four easiest color change magnitudes were offered during the easy perceptual blocks, while the four hardest color change magnitudes were offered during the hard perceptual blocks (two color change magnitudes appeared in both blocks). In other words, for Monkey Pu and Monkey Ra, the block shifted the color change magnitude frequency, while for Monkey Sa, the block shifted the color change magnitude range.

### Reward choice task

Before starting data collection on the change detection task, it was necessary to determine whether the animals understood the meaning of the reward cues. We recorded a single session in each animal of a two-choice task. After an initial fixation period, two movement targets with reward cues appeared in diametrically opposite positions in the periphery. Animals maintained fixation for a variable length options presentation period (500-900ms), then were given the opportunity to select either movement target. Animals were given ample time to decide and move and were immediately given the cued reward of the selected movement target upon a movement to one of the options. Choice preferences (Figure S6) indicated that the animals understood the reward cues.

#### Computing success rates

For each component of the task (overall, early aborts, perceptual, and motor), we computed success (or error) rates as a proportion pooled over trials from all recording sessions for each animal. Overall success rate was computed as (rewarded trials / total trials). Early abort ate was computed as (aborted trials / total trials). An aborted trial was defined as one in which the animal responded to the stimulus within the first 600ms of its presentation, a time during which no color change ever occurred. Detection performance (perceptual component of the task) was computed as (hits / non-aborted trials), where a hit was a trial in which the animal correctly detected the color change by responding within the window defined above. Motor performance was computed as (rewarded trials / hits). In Figure 1d-g, we reported the SE of the pooled proportion over trials from all recording sessions in each animal 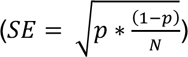, where *p* is the proportion and *N* is the number of trials in the corresponding proportion denominator pooled across recording sessions. Statistical significance was determined using a binomial test comparing the proportion on large reward trials and the proportion on jackpot trials.

#### Signal detection theory

Signal detection theory is traditionally applied to Go/No-go tasks to dissociate changes in perceptual sensitivity (d’) from changes in response bias (criterion). We adapted this framework to our task by treating each contrast cycle (every 240ms) of the stimulus as one Go/No-go opportunity. As such, each trial in our task contributed one or more Go/No-go opportunities depending on the length of the stimulus presentation period. Starting with the first possible color change contrast cycle (the third), for every subsequent contrast cycle until the color changed on that trial, we subdivided the animals’ behavioral responses into four categories: hit (responding within the response window when color change occurred), miss (not responding within the response window when color change occurred), false alarm (responding before the response window when color change did not occur), and correct rejection (not responding within the response window when color change did not occur). We then counted all hits, misses, false alarms, and correct rejections across all trials and contrast cycles. False alarms were attributed to the contrast cycle with the most recently closed response window.

For example, consider an example trial in which the color change occurred during the fifth contrast cycle (1080ms from stimulus onset). If the animal responded 250ms after the color change (within the acceptable response window defined in the “Motor component” section), this trial would contribute one hit for the fifth contrast cycle and two correct rejections for the third and fourth contrast cycles (the first and second contrast cycles are discarded, as no change can occur before the third). For that same example trial, if the animal responded only 10ms after the color change (before the response window for the fifth contrast cycle opened), the response would be considered a false alarm to the previous contrast cycle. In that case, the trial would contribute one false alarm for the fourth contrast cycle, and one correct rejection for the third contrast cycle.

From the counts of the four behavioral outcomes, we computed hit rate (hits / hits + misses), false alarm rate (false alarms / false alarms + correct rejections; Figure 1h), and miss rate (misses / hits + misses; Figure 1i). Perceptual sensitivity (d’) was defined as the difference in standard normal deviation between hit rate and false alarm rate (Equation 1; separation between distributions in schematic in Figure 1). Intuitively, this metric describes how well the animal can detect changes. Criterion (c) was defined as the negative average of the standard normal deviation of hit rate and false alarm rate (Equation 2; vertical line in schematic in Figure 1). Intuitively, criterion describes the threshold for responding to a color change, or a metric of the animal’s impulsivity where a higher criterion corresponds to a lower impulsivity. In each equation, *Z* indicates computation of the standard normal deviation using the inverse cumulative distribution function of a normal distribution, and *r* indicates a behavioral outcome rate.

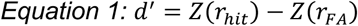

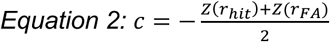

For Monkey Sa, who had a continuous stimulus presentation period distribution, we reassigned each trial to a stimulus presentation period bin, which spanned non-overlapping 200ms windows starting at the minimum (500ms) until the maximum (2500ms). After reassignment, we applied the same framework described above.

In contrast to the other success rate metrics described above, the metrics related to our signal detection theory analysis (false alarm rate, miss rate, d’, and criterion; Figure 1h-k) were computed per individual recording session. Statistical significance was thus determined by a paired t-test comparing the metrics for large reward trials and jackpot trials, paired across recording sessions.

#### Electrophysiological methods

Signals from the implanted electrode channels were band-pass filtered (0.3–7500 Hz) and then digitized at 30,000 Hz. For each channel, spiking waveforms were defined as a 52-sample (1.73ms) window of the filtered voltage signal triggered by the signal crossing a predefined threshold. The threshold was defined as a multiple (-4) of the root-mean-square voltage of a brief epoch of the raw signal on each channel collected at the beginning of the session. All behavioral and neural data were recorded using a Grapevine or TREK recording system (Ripple, Salt Lake City, UT) for further offline processing and analysis. All neural data in this work are from Monkey Pu and Monkey Ra in the Hand Task.

#### Preprocessing of neural data

We used a neural network to classify waveforms as spikes or noise. Details of the method were described previously in Issar, et al. 2020^91^. Briefly, a neural network was trained on human-sorted waveforms to distinguish between waveforms putatively of neural origin and waveforms not of neural origin. Using this network, we removed threshold crossings that were unlikely to be of neuronal origin and called the remaining waveforms “spikes”. The spikes from each channel on the Utah array were treated as multi-unit activity, and no further differentiation of single units was performed. Thus, each channel was treated as one neuron.

We excluded channels that had very low firing rates or very high trial-to-trial variability. To do this, we first binned neural activity on rewarded trials by counting spikes during the stimulus presentation period between stimulus onset and the end of the trial. We then removed channels with a mean spike rate lower than 2 spikes/s or a Fano factor greater than 10 spikes/s during this time window. This served to eliminate channels contaminated by recording artifacts, or channels that were not present for the entirety of a recording session. We also removed channels affected by artifactual crosstalk due to electrical coupling on the electrode array. For each pair of channels, we flagged their spikes as coincident if they occurred within 100μs of each other. If either channel in the pair had 20% or more of its spikes flagged as coincident, that pair was marked as having artifactual crosstalk. This procedure provided an index for each channel of how many other channels it was coincident with. We then iteratively removed the channel coincident with the highest number of other channels until no crosstalk remained, which ensured that the fewest possible number of channels were removed. After this process the number of remaining channels in Monkey Pu was 26.3 +/-6.8 (mean +/-standard deviation across sessions), and in Monkey Ra was 30.6 +/-8.2.

In Monkey Ra, there was one session where V4 was not recorded. This session was excluded from all neural analyses but remained in all behavioral analyses, yielding a total of 25 neural sessions for Monkey Ra. Additional session exclusion criteria such as minimum neuron or trial counts that were applicable to specific analyses are described in the corresponding analysis method.

#### Firing rate normalization

For the analyses in Figure 2c-d, firing rates of individual neurons were normalized by their peak firing rate to facilitate comparisons across neurons and recording sessions. Peak-normalization was performed for an individual neuron in the following manner. The maximum and minimum firing rate bin (10ms) during the first 600ms of the stimulus presentation period was identified from the average trace across all rewarded trials. The normalized firing rate (FR) was computed as (FR – FR_min_) / (FR_max_ –FR_min_), such that each neuron’s normalized firing rate fell between zero and one.

#### Spike count correlation

For the analysis in Figure 2e-f, we computed the mean spike count correlation (*r_sc_*) under each reward condition. We counted spikes in the time window 300-600ms from stimulus onset on rewarded trials in each reward condition. To match trial counts in each reward condition, we randomly subsampled trials without replacement to the minimum count 100 times. For each subsampled trial set, we computed the spike count correlation between each pair of neurons and averaged across all pairs of neurons for each experimental session. To get the mean *r_sc_* for each session, we averaged over the 100 subsamples in each reward condition.

#### Single-session factor analysis and %sv

We used factor analysis (FA)^95^ to identify shared trial-to-trial variability among the V4 population using latent variables. FA defines the linear-Gaussian relationship between spike count vectors (***x*** ∈ ℝ^3^) and latent variables (***z*** ∈ ℝ^4^) as:

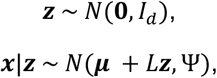

where *d* is the number of latent variables. The identity matrix *I* ∈ ℝ^4×4^ describes the prior covariance of each latent variable. For *m* neurons, *L* ∈ ℝ^3×4^ is the loading matrix that maps the neural activity into the *d* −dimensional latent space. The diagonal matrix Ψ ∈ ℝ^3×3^ contains the independent variance of each neuron, and the vector ***μ*** ∈ ℝ^3^ contains the mean spike counts of each neuron. FA estimates the sample covariance of the spike counts ***x*** as *C* = *LL*^6^ + Ψ using an exact expectation-maximization (EM) algorithm^96^.

An FA model was trained on spike counts from the first 600ms of the stimulus presentation period on a randomly subsampled (without replacement) 75% of the medium reward trials on each session. Sessions with less than 10 neurons were excluded, yielding 41 sessions for Monkey Pu (1 excluded) and 25 sessions for Monkey Ra (0 excluded). The dimensionality (number of latent variables, *d*) of the model was selected in a cross-validated manner. Candidate dimensionalities ranged from zero to ten (inclusive). From the 75% of medium reward trials, we then estimated the parameters of an FA model: the loading matrix *L*, and the private variance matrix Ψ. From these parameters, we computed the estimated covariance matrix as defined by the FA model framework (*C*).

For the analysis in Figure 2g-h, we took the remaining 25% of medium reward trials and all trials in the other reward conditions (small, large, and jackpot) and computed percent shared variance (%sv) in each reward condition separately. %sv indicates the proportion of each neuron’s spike count variance that was shared with other neurons in the population^52,94^. As the trials used to compute %sv were not used in the estimation of the FA model parameters, %sv is a cross-validated measure, and these trials were considered “held-out”. To compute held-out %sv, we first computed the sample covariance matrix (Σ = (*X* − ***μ***)(*X* − ***μ***)^6^) of each set of *N* held-out trials (*X* ∈ ℝ^3×(^, with mean ***μ*** ∈ ℝ^3^). The diagonal elements of Σ correspond to the total variances of each neuron. Held-out %sv was defined as the proportion of each neuron’s variance that was shared with other neurons. The denominator of this proportion was the corresponding diagonal element of Σ. The numerator of this proportion was computed as the corresponding diagonal element of the following matrix:

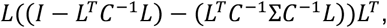

where *I* ∈ ℝ^4×4^ is a *d*-dimensional identity matrix.

In Figure 2g-h, we computed the held-out %sv on the remaining 25% of medium reward trials that were not used to train the FA model, and all trials in other reward conditions. To control for the trial count differences between reward conditions, we randomly subsampled trials without replacement from the held-out set in each reward condition to match the minimum number of trials across reward conditions. We repeated the subsampling procedure 100 times and averaged the held-out %sv across subsamples. This produced qualitatively similar results to those in Figure 2g-h (p=0.022, pooled across animals, n=66 sessions; Monkey Pu: mean Δ between jackpot and large = 0.89%, p=0.051, n=41; Monkey Ra: mean Δ = 0.61%, p=0.12, n=25; paired t-test).

#### Neural “stitching”

Because jackpot rewards occurred on only 5% of trials, to boost statistical power we employed a neural “stitching” algorithm that combined data across multiple recording sessions. We used a modified version of the neural stitching algorithm presented in Degenhart, et al.^56^, and described fully in Smoulder, et al.^28^. Briefly, the algorithm tracked channels that were stable across multiple recording sessions and leveraged this stability to identify a common latent space that all sessions mapped onto. This common latent space “stitched” the neural data across multiple sessions, significantly boosting trial counts for each analysis.

The first step in the stitching algorithm was the identification of stable channels. After the preprocessing described in the methods section “Preprocessing of neural data”, we identified channels that were stable across consecutive recording sessions, using the method described in Fraser & Schwartz^97^. Briefly, metrics such as firing rate and waveform shape were compared for a single channel on consecutive days. If the metrics were similar, the channel was noted as conserved across days. If not, the channel was noted as unique across days. Sessions at the beginning or end of the dataset with less than 10 conserved channels were removed to improve stability of the stitching model, yielding 33 sessions for Monkey Pu (9 excluded) and 23 for Monkey Ra (2 excluded). On average, the number of channels conserved across days was 14.6 for Monkey Pu and 17.2 for Monkey Ra.

The second step in the stitching algorithm was to identify a common latent space across recording sessions. After identifying conserved channels across day boundaries, we compiled conserved neural responses into an *M* × *T* matrix, where *M* was the total number of channels identified across all experimental sessions and *T* was the total number of selected time bins from trials across all experimental sessions. Elements of the matrix where a channel was not present for a certain session were left empty. To stitch the data across sessions, we identified a common latent space across all trials using FA (Figure S7). Specifically, the FA model was fit to the sample covariance matrix computed from the *T* samples, where some elements (where a channel was not present on that sample) were empty. To determine the dimensionality of the common latent space, we used five-fold cross validation to fit FA using the EM algorithm. We selected the dimensionality (*d*) that exceeded a threshold of 95% of the maximum cross-validated data likelihood from candidate dimensionalities ranging from one to ten (inclusive). The final FA stitching model was fit on these *d* dimensions using all data from all sessions (*d* = 9 dimensions for Monkey Pu, *d* = 8 for Monkey Ra). The original *M*-dimensional neural activity was projected into the *d*-dimensional latent space via a shared loading matrix (*L*^G^ ∈ ℝ^7×4^) identified by FA, effectively stitching it across multiple recording sessions. Here, the latent projections are referred to as neural activity or stitched activity for simplicity. Population average responses and reward modulation (Figure S8) in the stitched space remained qualitatively unchanged by this projection, indicating that the stitching procedure maintained the structure and trends present in V4 population responses from single sessions.

#### Computing the signal axis

To analyze color change signal in V4, we divided all trials in which the stimulus was correctly detected by the animal (hits) into two groups: Signal Present and Signal Absent. Signal Present trials were trials in which the color change happened during the earliest allowable contrast cycle (600ms from stimulus onset). Signal Absent trials were trials in which the color change happened later in the trial (>600ms), and therefore the signal was absent at 600ms.

To identify the dimension in V4 stitched activity that best captured the color change, we used linear discriminant analysis (LDA) to find an axis that separated stitched activity on Signal Present trials from Signal Absent trials. The Signal Present trials used for training were only those from color change magnitude 4 (the easiest color change). Stitched neural activity in 10ms bins ranging from 600ms to 800ms from stimulus onset (0 to 200ms from the color change in Signal Present trials) were used as training samples to fit a one-dimensional LDA axis. Stitched neural activity in 10ms bins ranging from 400ms to 800ms from stimulus onset (-200 to 200ms from the color change in Signal Present trials) was then projected onto the axis, and trial-averaged trajectories from all Signal Present (blue) and Signal Absent (gray) trials form the data shown in Figure 3b. Projected data for Signal Present trials included trials from all color change magnitudes. For signal axis projection on individual color change magnitudes, see Figure S5.

#### Selection of color change signal time window

To determine the time window over which to compute color change signal metrics, we fit a logistic regression model to predict the detection outcome (hit versus miss) from stitched neural activity near the color change. We tested window sizes of 10 to 120ms in increments of 10ms. We tested latencies ranging from 0ms after the color change to the minimum movement onset time (Monkey Pu: 180ms, Monkey Ra: 195ms). Trials with all stimulus presentation period lengths (not just Signal Present trials with the earliest color change time) were included in the regression training data. For each latency and window size tested, we randomly subsampled an equal number of hits and misses without replacement to match the trial count in each condition. We then performed 10-fold cross-validation to determine the cross-validated pseudo-*R*^2^ using McFadden’s method^98^. Across all window sizes and latencies, we identified the shortest time window at the earliest latency (relative to the color change) that achieved at least 95% of the maximum *R*^2^ from all tested configurations. For Monkey Pu, the resulting window was 20ms wide starting 80ms after the color change. For Monkey Ra, the window was 50ms wide starting 70ms after the color change.

#### Color change signal

We defined the color change signal as the difference in mean projection along the signal axis between Signal Present and Absent trials. For each Signal Present and Absent trial, we binned stitched activity in 10ms bins spanning the time window described above. We projected each bin onto the signal axis, then averaged the projections from each trial over the selected time window. This procedure yielded one average signal axis projection per trial, for Signal Present and Absent trials.

We split Signal Present and Absent trials by reward condition and computed the mean across trials per condition. To compare Signal Present and Absent trials, we took the difference (Δ) between the mean Signal Present and mean Signal Absent projection in each reward condition. Because we were subtracting two averages, to compute the SE of this quantity we used error propagation on the SE of each mean (Signal Present and Signal Absent; 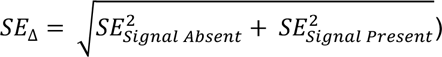. To determine significance, we performed a binomial proportion test where the proportion was the number of recording sessions where the average Δ was greater for large rewards than jackpots (Figure S5d).

#### Modulation depth

For the analysis in Figure 4b-c, we computed the movement target modulation depth for each neuron. All trials that were not aborted in the first 600ms after stimulus onset were included in this analysis (i.e., hits and perceptual error trials). Sessions with less than 5 trials in the minimum movement target-reward condition were excluded from this analysis, yielding 28 sessions for Monkey Pu (14 excluded) and 24 sessions for Monkey Ra (1 excluded). For each trial and each neuron (un-stitched), we computed its firing rate during the first 600ms of the stimulus presentation. Then, to match trial counts in each movement target-reward condition, we randomly subsampled trials to the count of the minimum condition. For each subsampled trial set, we computed the average firing rate across trials. Modulation depth was defined as the maximum minus the minimum firing rate across movement targets for each reward condition.

#### Movement target location decoding

For the analysis in Figure 4d-e, we decoded the location of the movement target under each reward condition in stitched neural activity. We trained an LDA decoder to predict movement target location (1 of 6) using a sliding window of overlapping 100ms bins of stitched activity during the stimulus presentation period. The bin centers were stepped by 10ms increments. To match trial counts in each reward condition, we randomly subsampled trials to the minimum count 100 times. For each of these hundred subsamples, we performed five-fold cross-validation, and each timepoint’s decoding accuracy was the average across all folds and repeats. The error on this estimate was computed as the SE of the average decoding accuracy, assuming a Bernoulli-distributed success rate. We repeated an analogous decoding analysis using neural activity from single sessions instead of stitched activity and found similar results (Figure S9).

