## Supplemental Information for "Eyes on the prize: a neural basis of perceptual choking under pressure"

#### Contents

1. Supplementary Fig. 1 with legend
2. Supplementary Fig. 2 with legend
3. Supplementary Fig. 3 with legend
4. Supplementary Fig. 4 with legend
5. Supplementary Fig. 5 with legend
6. Supplementary Fig. 6 with legend
7. Supplementary Fig. 7 with legend
8. Supplementary Fig. 8 with legend
9. Supplementary Fig. 9 with legend
10. Supplementary Table 1 with legend
11. Supplementary Table 2 with legend
12. Supplementary Information References

#### Supplementary Figure 1

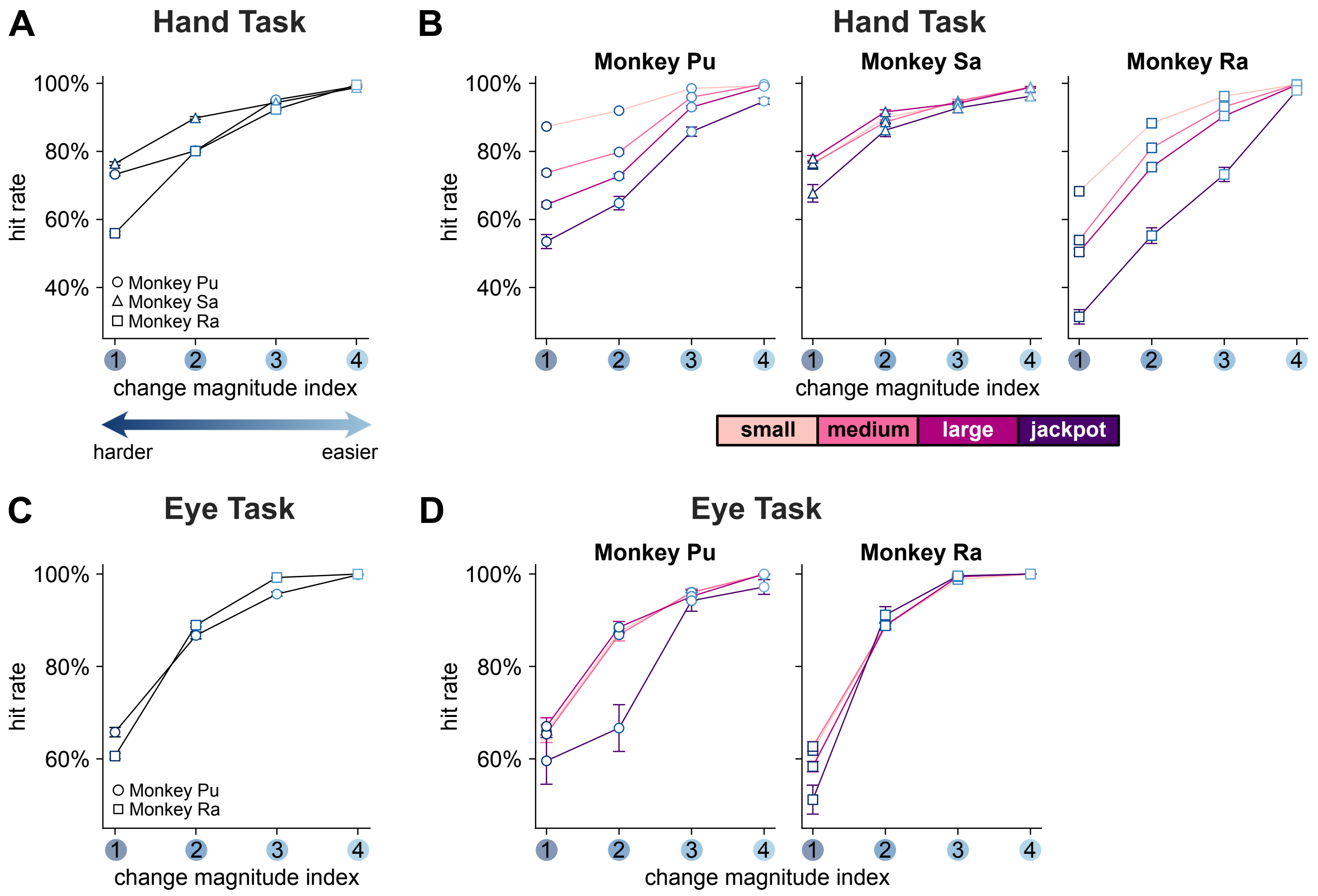


Figure S1: **Psychometric performance on Hand Task and Eye Task. Related to Figure 1.**

The perceptual component of the task was to detect a subtle color change. The difficulties of this color change were titrated for each animal such that it remained challenging. To understand how both perceptual difficulty and reward modulated detection behavior, here we report their performance as a function of both reward and perceptual difficulty (indexed from 1 to 4, hard to easy). Performance here is reported as hit rate (hits / hits + misses) as opposed to the metric in Figure 1f which was what we termed perceptual performance (hits / non-aborted trials). This change essentially removes false alarms from the denominator, which is relevant as false alarms should not be modulated by perceptual difficulty (by definition, the color never changed on a false alarm response). The horizontal axis position corresponds to the color change magnitude as in Figure 1c, where 1 is the hardest and 4 is the easiest. This forms a psychometric curve. Each plotted point indicates the mean $\pm$ standard error (SE) of hit rate across trials in a particular condition, computed assuming Bernoulli-distributed success rates. Note that in many cases the error bars are smaller than the plot symbol, obscuring them from view.

(A) First, we computed the hit rate across all trials in a given color change magnitude condition for each animal in the Hand Task (Monkey Pu: $n = [11108; 11444; 11757; 11697]$ total trials for color change magnitude $[1; 2; 3; 4]$, respectively, circles; Monkey Sa: $n = [6762; 6779; 6727; 6822]$, triangles; Monkey Ra: $n = [9587; 9521; 9689; 9775]$, squares). As expected, we found that performance increased as color change magnitude increased for all three animals.

(B) Next, to determine if both reward and perceptual difficulty modulated hit rates, we computed psychometric curves for trials in each reward condition (Monkey Pu, left; Monkey Sa, middle; Monkey Ra, right). The line color indicates the reward condition. Consistent with the increased miss rates (Figure 1i) and criterion (Figure 1k) we observed, hit rates tended to decrease with reward magnitude, especially for Monkey Pu and Monkey Ra. Together, these results show that hit rates increased as the perceptual difficulty became easier.

(C-D) Two animals (Monkey Pu and Monkey Ra) also performed a variant of the task in which they reported their detection with a saccadic eye movement to the movement target instead of a hand movement (Eye Task, see

Methods). The benefit of this task was that the motor demands were greatly simplified (saccade versus joystick movement), allowing us to better isolate perceptual behavior. Here, we assessed the animals' hit rates on the Eye Task by computing the same psychometric curves as above. Hit rates also increased with color change magnitude on this variant of the task (panel C; Monkey Pu: $n = [2151; 2054; 2051; 2034]$ trials per condition, circles; Monkey Ra: $n = [6236; 6054; 6195; 5912]$, squares). Hit rates were less modulated by reward on the Eye Task than on the Hand Task, but animals performed worse on jackpot trials (panel D, dark purple lines lower than the others; Monkey Pu, left; Monkey Ra, right). Independent of the motor effector (eyes or hands), hit rates increased with color change magnitude.

### Supplementary Figure 2

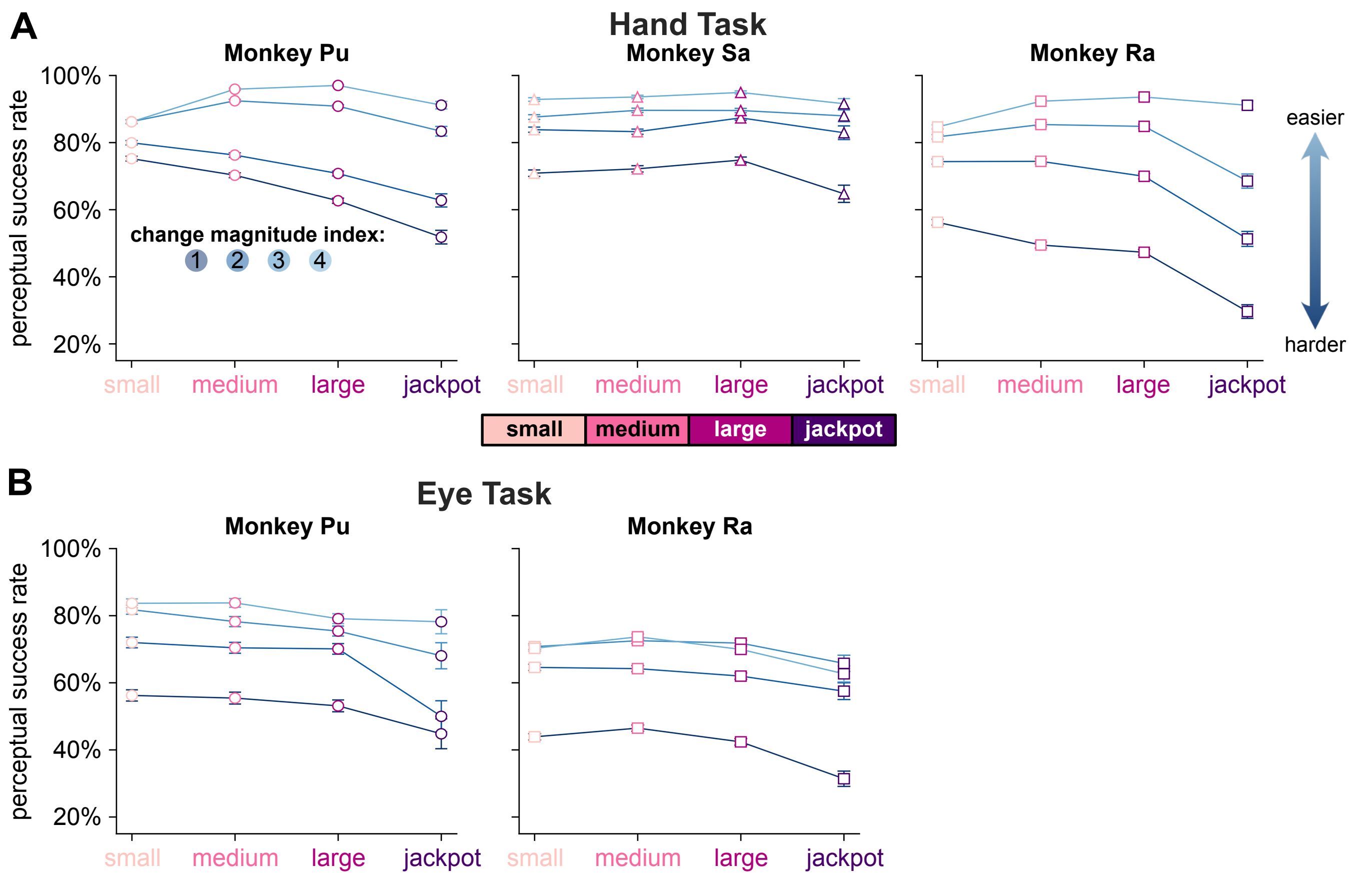


Figure S2: **Perceptual performance under each reward and change magnitude condition. Related to Figure 1.**

In Figure 1f, we reported the success rates on the perceptual component of the Hand Task. Here, we investigated whether the reward modulation of perceptual success rate was dependent on the difficulty of the color change. We computed the perceptual component success rate (hits / non-aborted trials) for each color change magnitude-reward condition separately. Each plotted line indicates the perceptual component success rate for a particular color change magnitude condition. The line color indicates the change magnitude index, where lighter colors are easier and darker colors are harder. Each point indicates the mean $\pm$ SE across trials in each condition. Note that in many cases the error bars are smaller than the plot symbol, obscuring them from view.

In the both the Hand Task (panel A) and the Eye Task (panel B) we found that reward modulated perceptual success rates under all color change magnitude conditions. In the Hand Task, perceptual performance significantly decreased on jackpot rewards compared to large rewards for Monkey Pu (left; $p < 0.0001$ for color change magnitude index 1, $p < 0.0001$ for index 2, $p < 0.0001$ for index 3, $p < 0.0001$ for index 4; binomial test), Monkey Sa (left; $p < 0.0001$ for index 1, $p = 0.011$ for index 2, $p = 0.18$ for index 3, $p = 0.0064$ for index 4; binomial test), and Monkey Ra (right; $p < 0.0001$ for index 1, $p < 0.0001$ for index 2, $p < 0.0001$ for index 3, $p = 0.017$ for index 4; binomial test). In the Eye Task, perceptual performance significantly decreased on jackpot rewards compared to large rewards for Monkey Pu (left; $p = 0.038$ for index 1, $p < 0.0001$ for index 2, $p = 0.028$ for index 3, $p = 0.43$ for index 4; binomial test) and Monkey Ra (right; $p < 0.0001$ for index 1, $p = 0.038$ for index 2, $p = 0.0058$ for index 3, $p = 0.00084$ for index 4; binomial test). It was not the case that perceptual choking under pressure only happened under easy or hard conditions, but was rather ubiquitous. Even so, the greatest performance decreases for jackpot trials tended to be for the harder change magnitude indices (darker lines). The average performance drop from large reward trials to jackpot trials across subjects for color change magnitude index 1 was $12.85\%$ for the Hand Task and $9.66\%$ for the Eye Task, while the corresponding drop in performance for index 4 was $3.90\%$ for the Hand Task and $4.10\%$ for the Eye Task. This further motivates why we titrated the color change magnitude values to be difficult.

### Supplementary Figure 3

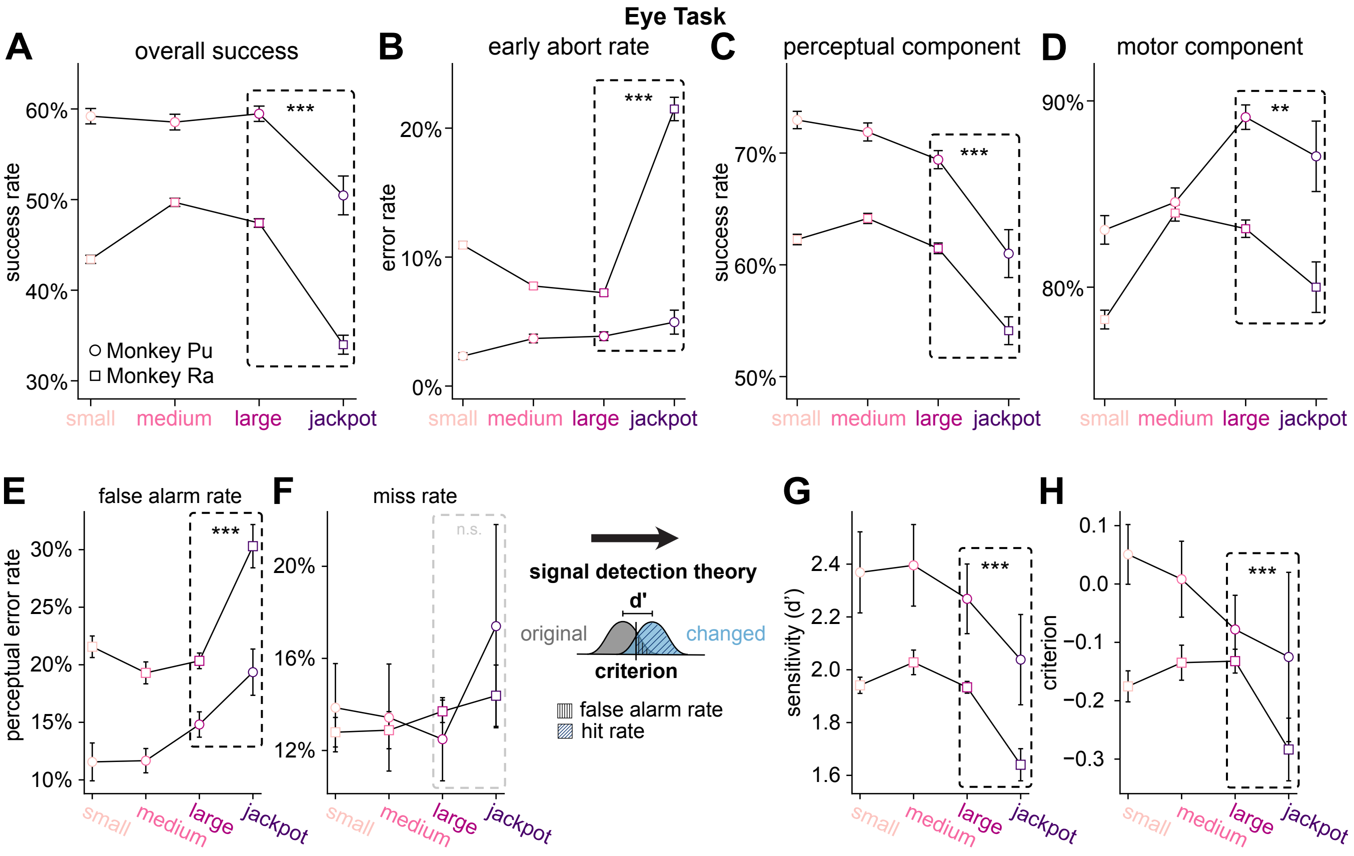


Figure S3: **Behavioral performance on Eye Task. Related to Figure 1.**

A subset of animals (Monkey Pu and Monkey Ra) performed two variants of the task: the Hand Task (Figure 1) in which they indicated their color change detection by moving a joystick-controlled cursor to the movement target, and the Eye Task in which they indicated with a saccadic eye movement. Here, we report the same behavioral metrics as in Figure 1, but for the Eye Task variant ($n = 31$ sessions total; Monkey Pu: $n = 10$; Monkey Ra: $n = 21$). As in Figure 1, each plotted point indicates the success rate $\pm$ SE across trials for panels A-D (computed assuming Bernoulli-distributed success rates) and across sessions for panels E-H. Note that in many cases the error bars are smaller than the plot symbol, obscuring them from view. Asterisks indicate significance on statistics pooled across animals (binomial test for panels A-D, and paired t-test for panels E-H). Three asterisks indicates a significance level of $p < 0.001$ and two asterisks indicates a significance level of $p < 0.01$.

(A-D) Overall (panel A), perceptual (panel C), and motor (panel D) performance in the Eye Task was consistent with the trends we observed in the Hand Task for both Monkey Pu (circles) and Monkey Ra (squares). Monkeys choked under pressure on their overall success rates ($p < 0.0001$, pooled across animals; Monkey Pu: $p < 0.0001$, Monkey Ra: $p < 0.0001$; binomial test), perceptual success rates ($p < 0.0001$, pooled across animals; Monkey Pu: $p < 0.0001$, Monkey Ra: $p < 0.0001$; binomial test), and motor success rates ($p = 0.0046$, pooled across animals; Monkey Pu: $p = 0.13$, Monkey Ra: $p = 0.0086$; binomial test). Though motor success rates trended downward on jackpot trials compared to large reward trials for both animals, the drop in performance was only significant for Monkey Ra. This was not necessarily unexpected, given that the motor demands for the Eye Task were not as difficult as in the Hand Task (i.e., an eye movement is less physically effortful than a hand movement). Early abort rates (panel B) increased on jackpot trials compared to large reward trials ($p < 0.0001$, pooled across animals; Monkey Pu: $p = 0.12$, Monkey Ra: $p < 0.0001$; binomial test), consistent with what we observed in the Hand Task.

(E-H) Reward modulation of the subtypes of perceptual errors also tended to be consistent with the Hand Task. Compared to large reward trials, on jackpot trials false alarm rates increased (panel E; $p < 0.0001$, pooled across animals, $n = 31$ sessions; Monkey Pu: $0.060$, $n = 10$; Monkey Ra: $p < 0.0001$, $n = 21$; paired t-test). Miss rates trended upwards, but the results were not significant (panel F; $p = 0.074$, pooled across animals, $n = 31$; Monkey Pu: $p = 0.064$, $n = 10$; Monkey Ra: $p = 0.32$, $n = 21$; paired t-test). We applied signal detection theory as shown in the schematic to assess changes to perceptual sensitivity (d') and criterion (see Methods). Similar to the Hand Task, on jackpot trials d' decreased compared to large reward trials (panel G; $p < 0.0001$, pooled across animals, $n = 31$; Monkey Pu: $p = 0.032$, $n = 10$; Monkey Ra: $p < 0.0001$, $n = 21$; paired t-test). In contrast to the Hand Task, criterion also decreased on jackpot trials (panel H; $p = 0.017$, pooled across animals, $n = 31$; Monkey Pu: $p = 0.35$, $n = 10$; Monkey Ra: $p = 0.0050$, $n = 21$; paired t-test). This could be due to the reduced motor demands on the Eye Task compared to the Hand Task. Even with this difference, there was still evidence for a distinct perceptual component of choking under pressure on this variant of the task.

#### Supplementary Figure 4

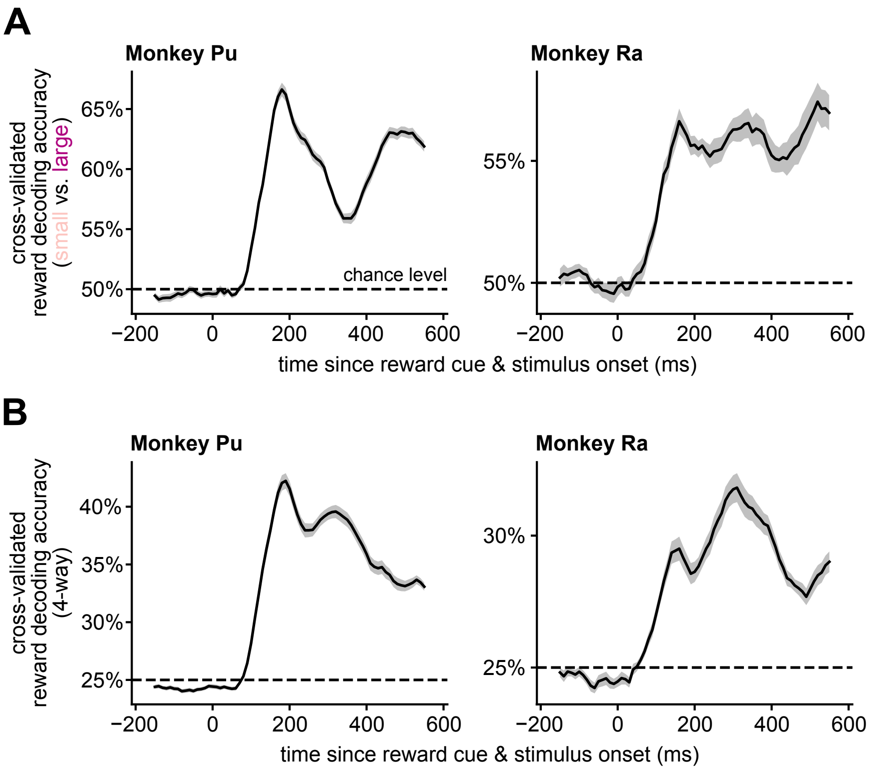


Figure S4: **Reward condition was decodable from V4 population activity. Related to Figure 2.**

In Figure 2, we found that, from small to large rewards, firing rates consistently increased among V4 neurons. When comparing jackpot trials to large reward trials, we also saw reliable modulation of firing rates, although inconsistent from animal to animal. For Monkey Pu, firing rates decreased for jackpots compared to large rewards while for Monkey Ra, they increased. To determine if the firing rate modulations by reward were statistically meaningful, we performed a decoding analysis. We trained a linear discriminant analysis (LDA) decoder to predict the reward condition from a sliding window of overlapping 100ms bins of neural activity during the stimulus presentation period. Sessions with less than ten neurons were excluded from this analysis, yielding $n = 41$ sessions for Monkey Pu (1 excluded) and $n = 24$ sessions for Monkey Ra (1 excluded). To avoid biasing the decoder, we balanced the number of trials in each reward condition by randomly subsampling to match the minimum count. Five-fold cross-validation was performed on the balanced counts to obtain the decoding accuracy. We repeated the subsampled cross-validation 100 times, and each session's decoding accuracy value is the average across all folds and repeats. The plotted shaded area indicates the mean $\pm$ SE of decoding accuracy across sessions. The horizontal axis position indicates the center of the 100ms bin of neural activity used to train the decoder.

(A) We performed two variants of this analysis. First, we investigated whether small versus large reward was differentiable in neural activity. To do this, we only used small and large reward trials to train the LDA decoder. We found that reward condition was decodable from V4 population activity in both Monkey Pu (left) and Monkey Ra (right). The chance level was $50\%$ (2 reward conditions, small and large).

(B) Next, we repeated the same analysis, but to decode all four reward conditions (small, medium, large, and jackpot). Even with more limited training samples due to the rarity of the jackpot trials, we found that reward condition was still decodable for both Monkey Pu (left) and Monkey Ra (right). Because all four reward conditions were included in this analysis, the chance level was $25\%$. Together, these results support that reward substantially modulated V4 activity and produced unique population activity states.

### Supplementary Figure 5

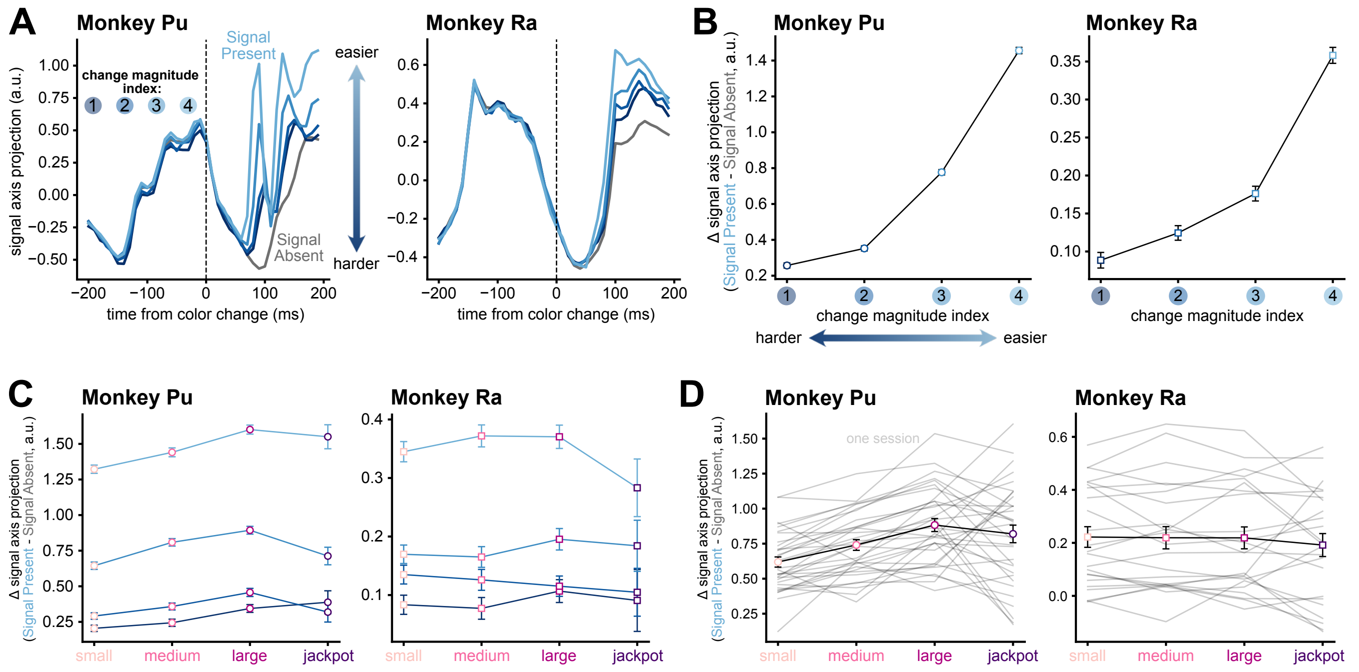


Figure S5: **The signal axis captured the color change signal in V4 population activity. Related to Figure 3.**

In Figure 3, we defined a signal axis, which captured V4 activity related to the color change (signal, see Methods). Here, we provide more details on how color change is represented along the signal axis.

(A) In Figure 3b, we visualized stitched V4 activity along the signal axis in the Signal Present and Signal Absent conditions. Here, we plotted the Signal Present trials in each color change magnitude condition separately, indicated by the line color, for Monkey Pu (left) and Monkey Ra (right). The individual change magnitude indices in the Signal Present condition range from darker (1, harder) to lighter blue (4, easier), as in Figure 1c. The Signal Absent condition is plotted in gray. Note, the Signal Absent condition for Monkey Ra (right) is identical to that shown in Figure 3b.

(B) To show how the different color change magnitudes fall on the signal axis, we computed the average $\Delta$ signal axis projection (Signal Present minus Signal Absent) within the same time window used in Figure 3c-d, but separated by color change magnitude index instead of reward condition. Here, we can see that the signal axis is indeed capturing the color change signal. From the hardest change (left of each panel) to the easiest change (right of each panel), the $\Delta$ signal axis projection increases. Intuitively, this indicates that V4 has more signal on the easier color changes. Each point indicates the mean $\Delta$ (mean of Signal Present condition minus the mean of the Signal Absent condition) $\pm$ SE (see Methods; Monkey Pu: left, Monkey Ra: right). Note that in many cases the error bars are smaller than the plot symbol, obscuring them from view.

(C) In panel B, we saw that the color change magnitude has a dramatic effect on the magnitude of the $\Delta$ signal axis projection. In Figure 3c-d, we found that reward also impacts $\Delta$ signal axis projection. Here, we tested whether the trends elicited by the reward condition were consistent within each individual color change magnitude. We computed the average $\Delta$ signal axis projection in each color change magnitude-reward condition. The $\Delta$ signal axis projections tended to decrease on jackpot trials compared to large reward trials across all color change magnitudes for both Monkey Pu (left) and Monkey Ra (right). Similar to how perceptual behavior was decreased on jackpots in all color change magnitude conditions (Figure S2), the neural correlate of color change was also reduced on jackpots in almost all conditions.

(D) To compute statistics on the $\Delta$ signal axis projections in Figure 3c-d, we considered trials in each reward condition further subdivided by the recording session. Each gray trace is the average $\Delta$ signal axis projection computed from trials in a single recording session. The black traces indicate the mean $\pm$ SE across sessions for Monkey Pu (left) and Monkey Ra (right). Individual session values for large and jackpot rewards were used to compute the binomial statistics in Figure 3c-d. Across individual sessions, we found that the $\Delta$ signal axis projections tended to decrease on jackpot trials compared to large reward trials for Monkey Pu ($n = 21/33$ sessions greater for large reward, $p = 0.081$) and Monkey Ra ($n = 14/23$ sessions greater for large reward, $p = 0.20$). While neither animal's individual sessions reached significance, pooling across sessions from both animals supported the claim that the color change signals are reduced on jackpot trials compared to large reward trials ($n = 35/56$, $p = 0.041$).

### Supplementary Figure 6

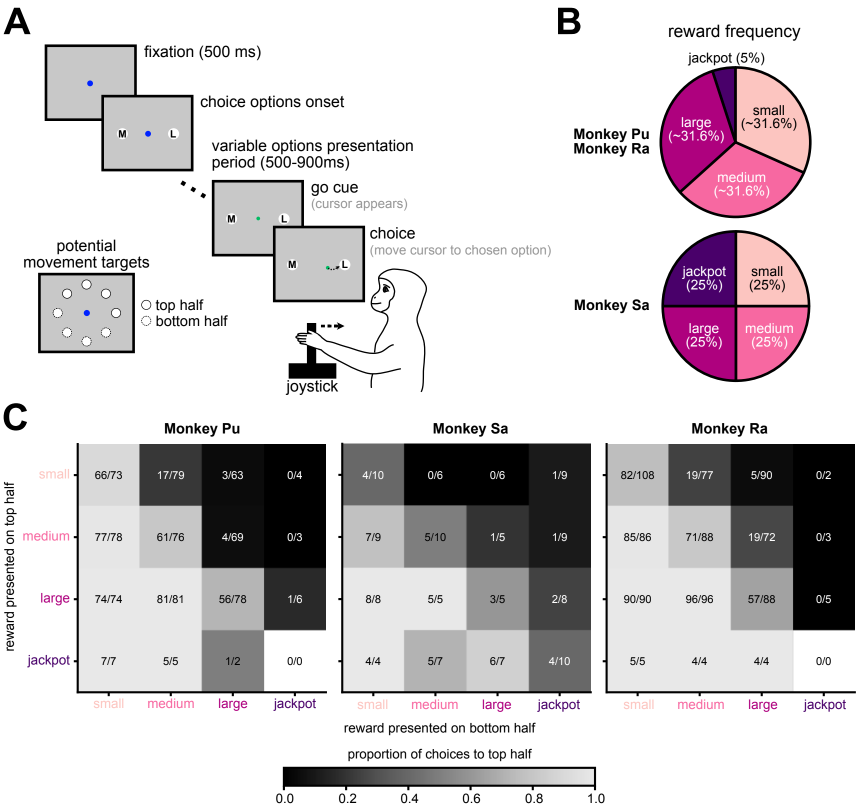


Figure S6: **Behavioral performance on Choice Task. Related to Methods.**

(A) To determine if animals understood the meaning of the reward cues before we recorded the data used in this work, we ran one session of a reward choice task. The choice task was structured similarly to the main Hand Task (see Methods), but two movement target options with a reward cue were offered on every trial, in diametrically opposed locations. After a variable options presentation period (500-900ms on each trial), animals were given a go cue to freely choose one of the two reward options with ample time. Animals selected by moving a joystick-controlled cursor (green circle) to one of the two options. Upon reaching one of the movement targets, animals received the reward that was cued on the selected movement target.

(B) Rewards were offered at the same frequency as in the main Hand Task on every movement target (two per trial) for Monkey Ra and Monkey Pu. For Monkey Sa, each reward, including jackpot, was offered equally frequently. For Monkey Ra and Monkey Pu, an additional constraint was applied such that two jackpots did not appear in the same trial.

(C) To quantify the animals' choice behavior, we constructed confusion matrices for each animal (Monkey Pu: left, Monkey Sa: middle, Monkey Ra: right). The rows indicate the reward presented on the top half of the screen (options shown with solid outlines in the inset of panel A). The columns indicate the reward presented diametrically opposite on the bottom half of the screen (options shown with dashed outlines in the inset of panel A). The color of each square indicates the proportion of choices made under each condition, with the number of trials displayed on the square.

The animals made logical choices (choosing the larger of the two reward offers) on the majority of trials (Monkey Pu: $94.27\%$ of choices between unequal options were logical, Monkey Sa: $87.95\%$, Monkey Ra: $91.76\%$). Elements above the diagonal were darker colored, indicating that the animal more often chose the reward on the bottom half of the screen, which in these conditions was greater than the one on the top half. Elements below the diagonal were lighter colored, indicating that the animal more often chose the reward on the top half of the screen, which in these conditions was greater than the one on the bottom half. The diagonal elements indicate conditions in which an equal reward was presented in the two locations, and the proportions here served to indicate spatial bias.

#### Supplementary Figure 7

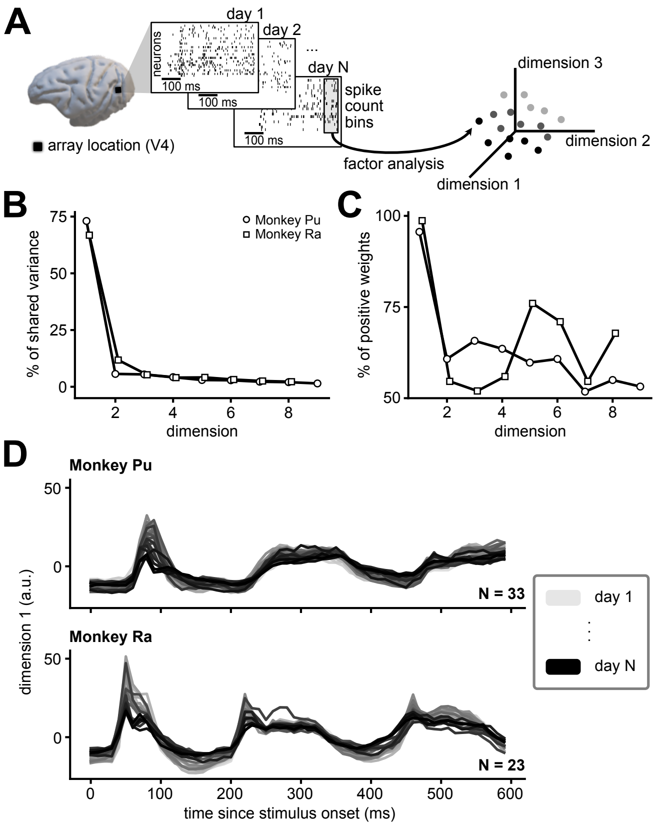


Figure S7: **Stitching model metrics. Related to Methods.**

Because jackpot rewards occurred on only 5% of trials, to boost statistical power we employed a neural stitching algorithm that combined data across multiple recording sessions. To do this, we adapted the stitching procedure used in our previous work in motor cortex [1], and initially presented in Degengart, et al. [2] (see Methods). Briefly, we first identified recording channels that were present and stable across multiple recording sessions [3]. Then, a factor analysis (FA) [4] model was trained using this tracked neural activity across all sessions. We refer to this procedure as "stitching" the neural activity together across recording sessions, and the resulting latent space as the "stitched" space.

(A) The result of the neural stitching algorithm is the ability to combine data from each recording session (each raster plot, left) into a common latent space identified by FA (right). Each spike count bin from each recording session (e.g., in the shaded box, left) was projected into the latent space and formed one element of our "stitched" data.

(B) To understand how the stitching procedure captured features of our V4 recordings, we characterized a few statistics of the FA model. The stitching procedure identified a 9-dimensional latent space for Monkey Pu and an 8-dimensional latent space for Monkey Ra. First, we assessed the relative strength of each dimension, by computing the proportion of the population shared variance that was explained by each dimension. The dimensions were ordered by the amount of variance explained, such that dimension 1 explained the most, dimension 2 explained the second-most, etc. The percentage of shared variance explained by the $i^{\text{th}}$ dimension of the FA model was computed as $\frac{\lambda_i}{\text{tr}(\tilde{L}\tilde{L}^T)}$, where $\lambda_i$ is the $i^{\text{th}}$ eigenvalue of $\tilde{L}\tilde{L}^T$, $\tilde{L} \in \mathbb{R}^{M \times d}$ is the loading matrix mapping $M$ neurons to the $d-$dimensional stitched latent space, and $\text{tr}(\cdot)$ indicates the trace (see Methods for full definitions of stitching model parameters). Across both animals (Monkey Pu: circles, Monkey Ra: squares), we found that dimension 1 explained an overwhelming proportion of the shared variance in our V4 recordings relative to the rest of the dimensions (sharp decline from dimension 1 to dimension 2). This indicated that much of the shared variance could be attributed to a single mode of variance.

(C) We next assessed the extent to which the latent variables identified by FA reflected the neurons all increasing or decreasing their activity together by looking at the diversity of weights onto the population of neurons. To quantify this, we computed the percentage of neurons that shared a similar sign along each dimension of the stitching model. We oriented the sign of each dimension such that the majority of weights were positive (i.e., the minimum percentage of same-signed weights is 50%, which we notate as positive by convention). This sign change was only performed for this analysis. In both animals (Monkey Pu: circles, Monkey Ra: squares), dimension 1 was comprised of weights that were almost 100% positive, indicating that it was very tightly linked to changes in firing rates among the neurons. In other words, an increase in activity along the first dimension of the stitched space indicated a general increase in firing of the underlying neural population. The other dimensions had a higher diversity of weights onto the neurons (50-75% positive).

(D) Because we found that dimension 1 was tightly linked to the firing rates of the population, we used this as a baseline to assess how stable the stitched trial-averaged response was across recording sessions. Though the stitching procedure should seamlessly track neural activity across sessions, it is possible that a mismatch of dimensions could occur between sessions. We visualized stitched activity along dimension 1 separately for each recording session (one line per session) for Monkey Pu (top, $n = 33$ sessions) and Monkey Ra (bottom, $n = 23$ sessions). Darker lines indicate sessions near the end of each dataset, and lighter colors indicate sessions near the beginning of each dataset. Stitched activity from all sessions were heavily overlapping with no sudden changes from session to session, indicating that the stitching procedure preserved neural activity patterns across sessions.

### Supplementary Figure 8

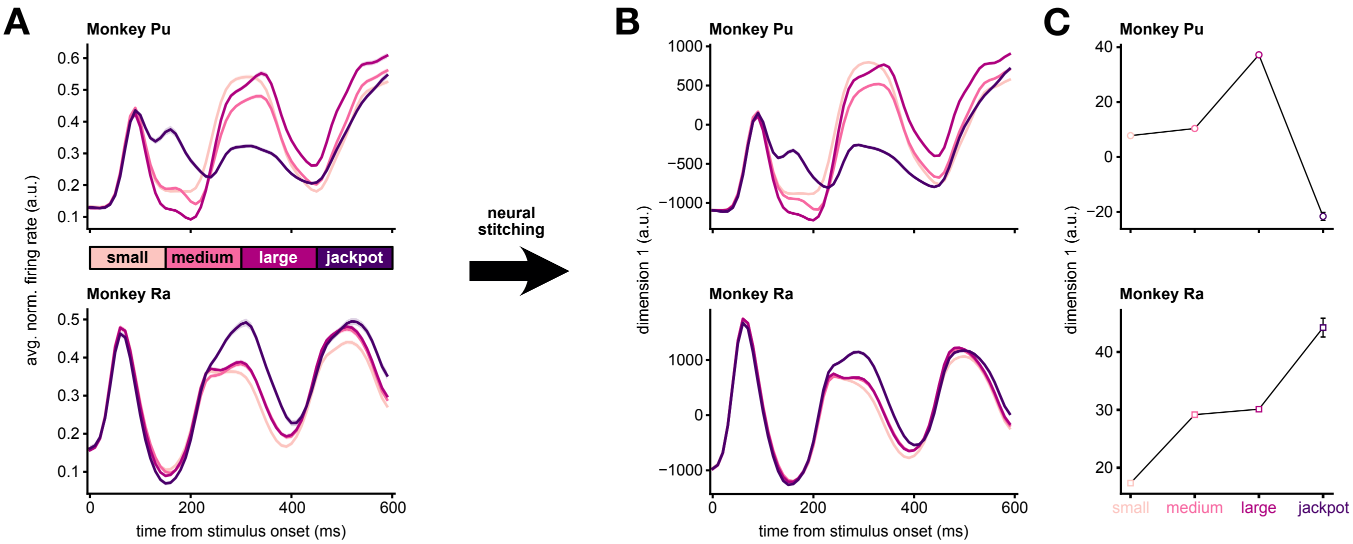


Figure S8: **Stitched space preserved reward modulation. Related to Methods.**

The neural stitching procedure combined neural activity across all sessions. As shown in Figure S7d, the stitching procedure reflected a consistent response along the first dimension across days. To validate the robustness of the stitching procedure, here we compared whether reward trends were affected by the stitching procedure.

(A) As a point of comparison, we computed the population average response under each reward condition in single-session neural activity (i.e., the un-stitched data). Activity from each neuron and each session was peak-normalized (see Methods), then the normalized neurons were combined across sessions. We averaged across trials in each reward condition over all neurons. For visualization purposes, we smoothed the average trace using a Gaussian kernel with a width of 15ms. The plotted shaded area indicates the smoothed mean $\pm$ SE across neurons for Monkey Pu (top, $n = 1105$ neurons) and Monkey Ra (bottom, $n = 796$ neurons). Note that the shaded area may be smaller than the plotted line, obscuring it from view. The population average response exhibited reward tuning throughout the stimulus presentation period. These reward modulation trends can now serve as the baseline to compare against the stitched activity (dimension 1). As dimension 1 in the stitched space had an overwhelming majority of positive-signed weights (Figure S7b), the population average is a particularly relevant comparison.

(B) After performing the neural stitching procedure, we computed the average activity along dimension 1 for each reward condition separately. For visualization purposes, we smoothed the average trace using a Gaussian kernel with a width of 15ms. The plotted shaded area indicates the smoothed mean $\pm$ SE across trials in each condition for Monkey Pu (top; $n = 9782$ small, $n = 10523$ medium, $n = 10056$ large, and $n = 1473$ jackpot trials) and Monkey Ra (bottom; $n = 9170$ small, $n = 8836$ medium, $n = 8793$ large, and $n = 1089$ jackpot trials). Note that the shaded area may be smaller than the plotted line, obscuring it from view. We found that the reward trends were qualitatively consistent with the population average firing rate, indicating that the stitching procedure preserved reward modulation.

(C) To further emphasize this consistency, we performed an analogous analysis to that in Figure 2d. We computed the average activity along dimension 1 late in the stimulus presentation period (300-600ms after stimulus onset, same as Figure 2d) on trials in which the animal would go on to correctly detect the color change. Compared to Figure 2d, the reward trends in each animal (Monkey Pu, top; Monkey Ra, bottom) were similar. Each point indicates the mean $\pm$ SE across trials in each condition (same $n$ values as in panel B). Note that in many cases the error bars are smaller than the plot symbol, obscuring them from view.

### Supplementary Figure 9

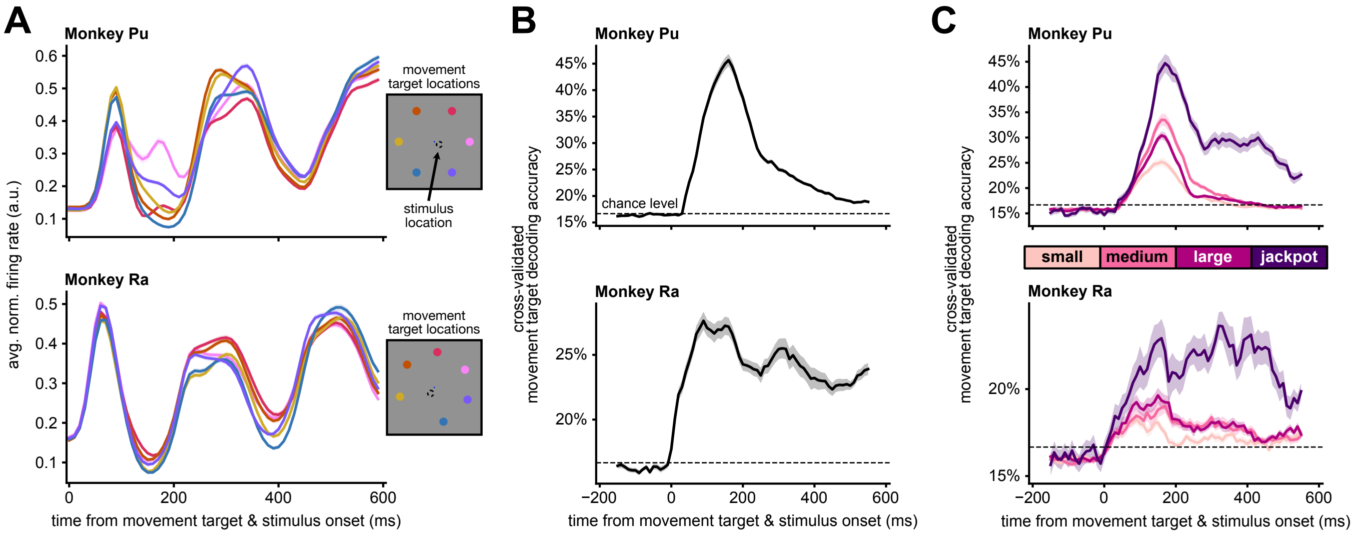


Figure S9: **Movement target decoding in single sessions. Related to Figure 4.**

In Figure 4, we decoded the movement target location from stitched V4 activity. The motivation for investigating this question in stitched activity (rather than single sessions) was to avoid trial limitations due to the jackpots being rare. Here, we investigated whether an effect consistent with that in Figure 4d-e could be found on single sessions. More specifically, whether movement target decoding accuracy on jackpot trials would be greater than for other rewards and sustained throughout the stimulus presentation period.

(A) To determine whether differences in single-session neural activity could be seen in firing rates, we first computed the population average response under each movement target condition in the single-session data. In a similar analysis to Figure S8a, we peak-normalized the activity of each individual neuron then combined across neurons (see Methods). We trial-averaged activity by movement target condition and averaged across all neurons. For visualization purposes, we smoothed the average trace using a Gaussian kernel with a width of 15ms. The plotted shaded area indicates the smoothed mean ± SE across neurons for Monkey Pu (top, n = 1105 neurons) and Monkey Ra (bottom, n = 796 neurons). Note that the shaded area may be smaller than the plotted line, obscuring it from view. The line color indicates the movement target condition, the location of which is shown on the inset to the right of each panel. The population average response exhibited movement target tuning throughout the stimulus presentation period. This indicated that movement target may be able to be decoded on a single-session basis.

(B-C) To decode the movement target location on single sessions, we used linear discriminant analysis (LDA) to separate single-session (un-stitched) neural activity under different movement target conditions. We trained a separate LDA decoder using a sliding window of overlapping 100ms bins of V4 population spike count vectors. The horizontal axis position indicates the center of the 100ms bin of neural activity used to train the decoder. Plotted shaded area indicates the mean ± SE of decoding accuracy across sessions (Monkey Pu: $n = 42$, Monkey Ra: $n = 25$). The chance level is $\approx 16.6\%$ (6 movement target conditions).

(B) We first investigated whether the movement target location was at all decodable on a single-session basis. Without subdividing by reward (as in Figure 4), we trained an LDA decoder to predict movement target location and found that we could predict the location well above chance level beginning shortly after movement target and stimulus onset ($t = 0$ms) for both Monkey Pu (top) and Monkey Ra (bottom). For each session, we performed 5-fold cross-validation to compute decoding accuracy, which was defined as the average across individual folds.

(C) After finding that we could reliably decode the movement target location on a single-session basis, we next tested whether the reward modulation we observed in the stitched neural activity (Figure 4) was also evident under the trial limitations of single sessions. Sessions without a minimum of 6 trials in each movement target-reward condition were excluded from this analysis, yielding $n = 26$ sessions for Monkey Pu (16 excluded) and $n = 21$ sessions for Monkey Ra (4 excluded). We performed a similar analysis as in Figure 4 (see Methods), but using 100ms bins of single-session V4 spike count vectors instead of stitched activity. As in Figure 4, we randomly subsampled trials in each movement target-reward condition to match the minimum count across all movement target-reward conditions. Because of trial limitations, cross-validation was performed using leave-one-out, rather than k-fold. Even when limited by trial counts of single sessions, we found reward modulation consistent with what we observed in the stitched activity. On jackpot trials, movement target decoding accuracy was both greater than for other rewards and sustained throughout the stimulus presentation period in both Monkey Pu (top) and Monkey Ra (bottom).

### Supplementary Table 1

Table S1:

| Monkey Name | Task Name | # of Sessions | Small | Medium | Large | Jackpot |
|---|---|---|---|---|---|---|
| Monkey Sa | Hand Task | 3 | 0.05 | 0.15 | 0.4 | 2 |
| Monkey Sa | Hand Task | 22 | 0.057 | 0.171 | 0.456 | 2.28 |
| Monkey Pu | Hand Task | 42 | 0.057 | 0.171 | 0.456 | 2.28 |
| Monkey Ra | Hand Task | 6 | 0.072 | 0.156 | 0.232 | 1.52 |
| Monkey Ra | Hand Task | 1 | 0.049 | 0.156 | 0.232 | 1.52 |
| Monkey Ra | Hand Task | 5 | 0.042 | 0.148 | 0.224 | 1.52 |
| Monkey Ra | Hand Task | 5 | 0.034 | 0.148 | 0.224 | 1.52 |
| Monkey Ra | Hand Task | 9 | 0.032 | 0.133 | 0.209 | 1.52 |
| Monkey Pu | Eye Task | 1 | 0.057 | 0.114 | 0.228 | 1.52 |
| Monkey Pu | Eye Task | 9 | 0.057 | 0.171 | 0.456 | 2.28 |
| Monkey Ra | Eye Task | 11.5 | 0.095 | 0.19 | 0.285 | 1.9 |
| Monkey Ra | Eye Task | 1.5 | 0.076 | 0.152 | 0.228 | 1.52 |
| Monkey Ra | Eye Task | 4 | 0.053 | 0.160 | 0.266 | 1.52 |
| Monkey Ra | Eye Task | 5 | 0.053 | 0.160 | 0.213 | 1.52 |

**Table S1: Reward volumes in milliliters. Related to Methods.**

For each animal, we titrated the volume of reward to motivate performance. Volumes scaled up by approximately 1x, 3x, 8x for small, medium, and large reward. Jackpot rewards were much larger ($\approx$40x). Here, we report the volume of each reward condition in milliliters, as well as the number of sessions collected under each configuration. A value of 0.5 indicates that the reward volumes were adjusted during a particular recording session to better motivate the animal.

#### Supplementary Table 2

Table S2:

| Monkey Name | Task Name | # of Sessions | Color change magnitude index | | | | | |
|---|---|---|---|---|---|---|---|---|
| | | | 1 | 2 | 3 | 4 | 5 | 6 |
| Monkey Sa | Hand Task | 17 | 5.5 | 5.75 | 6 | 6.75 | 13 | 20 |
| Monkey Sa | Hand Task | 2 | 5.5 | 6 | 6.25 | 7.25 | 13 | 20 |
| Monkey Sa | Hand Task | 1 | 5.5 | 6 | 6.5 | 7.5 | 13 | 20 |
| Monkey Sa | Hand Task | 1 | 6 | 6.5 | 7 | 7.5 | 17 | 20 |
| Monkey Sa | Hand Task | 4 | 6 | 6.5 | 7 | 11 | 17 | 20 |
| Monkey Pu | Hand Task | 6 | 7 | 7.5 | 10 | 14 | | |
| Monkey Pu | Hand Task | 4 | 7 | 8 | 11 | 18 | | |
| Monkey Pu | Hand Task | 32 | 8 | 9 | 12 | 20 | | |
| Monkey Ra | Hand Task | 1 | 4 | 5.5 | 7 | 12 | | |
| Monkey Ra | Hand Task | 3 | 4 | 5.5 | 7 | 15 | | |
| Monkey Ra | Hand Task | 2 | 4.5 | 6 | 7.5 | 15 | | |
| Monkey Ra | Hand Task | 5 | 5 | 6.5 | 8 | 12 | | |
| Monkey Ra | Hand Task | 1 | 5 | 6.5 | 8 | 13 | | |
| Monkey Ra | Hand Task | 5 | 5 | 6.5 | 8 | 16 | | |
| Monkey Ra | Hand Task | 1 | 6 | 7 | 9 | 15 | | |
| Monkey Ra | Hand Task | 1 | 6 | 8 | 11 | 18 | | |
| Monkey Ra | Hand Task | 1 | 6 | 9 | 12 | 18 | | |
| Monkey Ra | Hand Task | 4 | 7 | 11 | 14 | 21 | | |
| Monkey Ra | Hand Task | 2 | 9 | 12 | 15 | 22 | | |
| Monkey Pu | Eye Task | 5 | 3 | 4 | 5 | 10 | | |
| Monkey Pu | Eye Task | 2 | 3.5 | 5 | 6 | 10 | | |
| Monkey Pu | Eye Task | 1 | 4 | 5.5 | 7 | 12 | | |
| Monkey Pu | Eye Task | 1 | 5 | 6.5 | 8 | 13 | | |
| Monkey Pu | Eye Task | 1 | 6 | 8 | 10 | 15 | | |
| Monkey Ra | Eye Task | 21 | 3 | 5 | 8 | 13 | | |
| Monkey Ra | Eye Task | 0.5 | 4 | 6 | 9 | 14 | | |
| Monkey Ra | Eye Task | 0.5 | 6 | 9 | 12 | 18 | | |

**Table S2: Color change magnitudes. Related to Methods.**

For each animal, we titrated the color change magnitudes across sessions to prevent them from reaching their detection performance ceiling. As the animals gained more experience with a particular color change magnitude, their detection performance improved. We gradually decreased the color change magnitudes (increased difficulty) over many sessions to compensate for their improved performance. The color change was defined as a shift in hue space, measured in degrees. We report the color change magnitudes in degrees, as well as the number of sessions collected under each configuration. For Monkey Sa, color change magnitudes were one of six graded difficulties on each trial, which are listed in the named columns 1-6 in the table. For Monkey Pu and Monkey Ra, color change magnitudes were one of four graded difficulties, which are listed in the named columns 1-4 in the table. A value of 0.5 indicates that the color change magnitude values were adjusted during a particular recording session to better capture the range of psychometric performance (as in Figure S1).
